# Quantitative systems pharmacology modeling of pyrrolobenzodiazepine antibody-drug conjugates targeting BCMA

**DOI:** 10.1101/2025.02.20.639376

**Authors:** Inez Lam, David Dai, Rosalinda H. Arends, Venkatesh Pilla Reddy, Kathryn Ball, Feilim Mac Gabhann

## Abstract

Antibody-drug conjugates (ADCs) are novel therapeutics combining two molecules linked together: an antibody that provides targeting to specific cells, and a cytotoxic drug (warhead) that can deconjugate from the antibody and kill those cells. The warhead by itself would be too nonspecific and toxic; the antibody by itself would be insufficiently effective at killing the cells. As more and more ADCs enter the drug development pipeline, understanding the mechanistic reasons behind efficacy and toxicity is critical to evaluating both successful and failed trials. Here, we have developed a mechanistic computational model of ADCs, and specifically parameterize it using data for MEDI2228, an anti-BCMA antibody conjugated to pyrrolobenzodiazepine (PBD) warheads. We build the model to track not only the concentrations of ADC and its released warhead drug inside and outside the cell, but also to track the recent history of the warhead, so that we can distinguish between pathways for on-target and bystander (nontargeted) cell killing. We show that this effect is predicted to be small under *in vitro* conditions due to dilution in large volumes of media, but likely to form a significant part of both targeted and nontargeted cell killing *in vivo*, where the extracellular volume has less of a dilution effect. We also explore the impact of key design parameters of ADCs, including drug to antibody ratio (DAR), warhead potency, and lipophilicity; this analysis demonstrates the balance needed between killing of targeted and nontargeted cells. Using this quantitative systems pharmacology model, we can generate insights for optimization of ADC design and determine which factors are most critical to efficacy and toxicity, leading to more informed and rational development of cancer therapies.

## Introduction

In the quest for effective cancer treatment, one of the biggest challenges in therapy design is creating a wide therapeutic window, i.e. designing the therapy such that a wide range of drug concentrations or doses are clinically useful in leading to therapeutic response. With a narrow therapeutic window, both underdosing (ineffective) and overdosing (potentially toxic) become likely. A wide therapeutic window maximizes efficacy while minimizing toxicity across a wide range of drug dosing. Although traditional chemotherapy can provide the desired cell-killing capabilities, the lack of discrimination between cancerous and healthy cells can cause toxic side effects [1,2]. Some of these side effects can be mitigated by supportive care [3,4], but other side effects can be debilitating or worse, so minimizing toxicity is still a key goal in the development of anticancer therapeutics [5–8]. In contrast to the more indiscriminate killing of cells by chemotherapy, antibodies can be highly targeted and bind to specific antigens on the surface of the target cells [9,10]. However, while some antibodies can mediate cell killing in some cancers, by themselves they may lack the desired level of anti-tumor or cell-killing capabilities in other cancers [11–13]. Primarily developed as cytotoxic cancer therapies, antibody-drug conjugates (ADCs) are designed to harness the cytotoxicity of chemotherapy and the specificity of antibodies, while limiting their disadvantages [14,15].

ADCs are engineered immunoconjugate drugs comprising 3 core components: (1) a monoclonal antibody (Ab) with (2) one or more cytotoxic small molecules (known as warheads or payloads), attached via (3) a chemical linker (**Figure 1A**). The antibody targets a specific cancer-associated antigen (which can be different for different cancer types), so the warhead can selectively or preferentially kill cancer cells while sparing most healthy tissue. This modular structure with multiple components means that ADCs have multiple design levers with which to optimize efficacy and minimize toxicity [14,16–18]. Thus, ADC design requires a deep understanding of the underlying biological and pharmacological mechanisms. While experimental data characterizing ADCs is essential, using experimental methods alone to acquire this understanding can be laborious, expensive, or even infeasible. Quantitative systems pharmacology (QSP) modeling can combine mechanistic knowledge with available preclinical and clinical data to make predictions and gain insights through quantitative simulation of drug action and performance [19,20]. Use of QSP approaches to construct interpretable and predictive models has been increasing in recent years, particularly by biopharmaceutical companies to support decision-making in drug development, drug approvals, and clinical studies [21,22].

**Figure 1.**
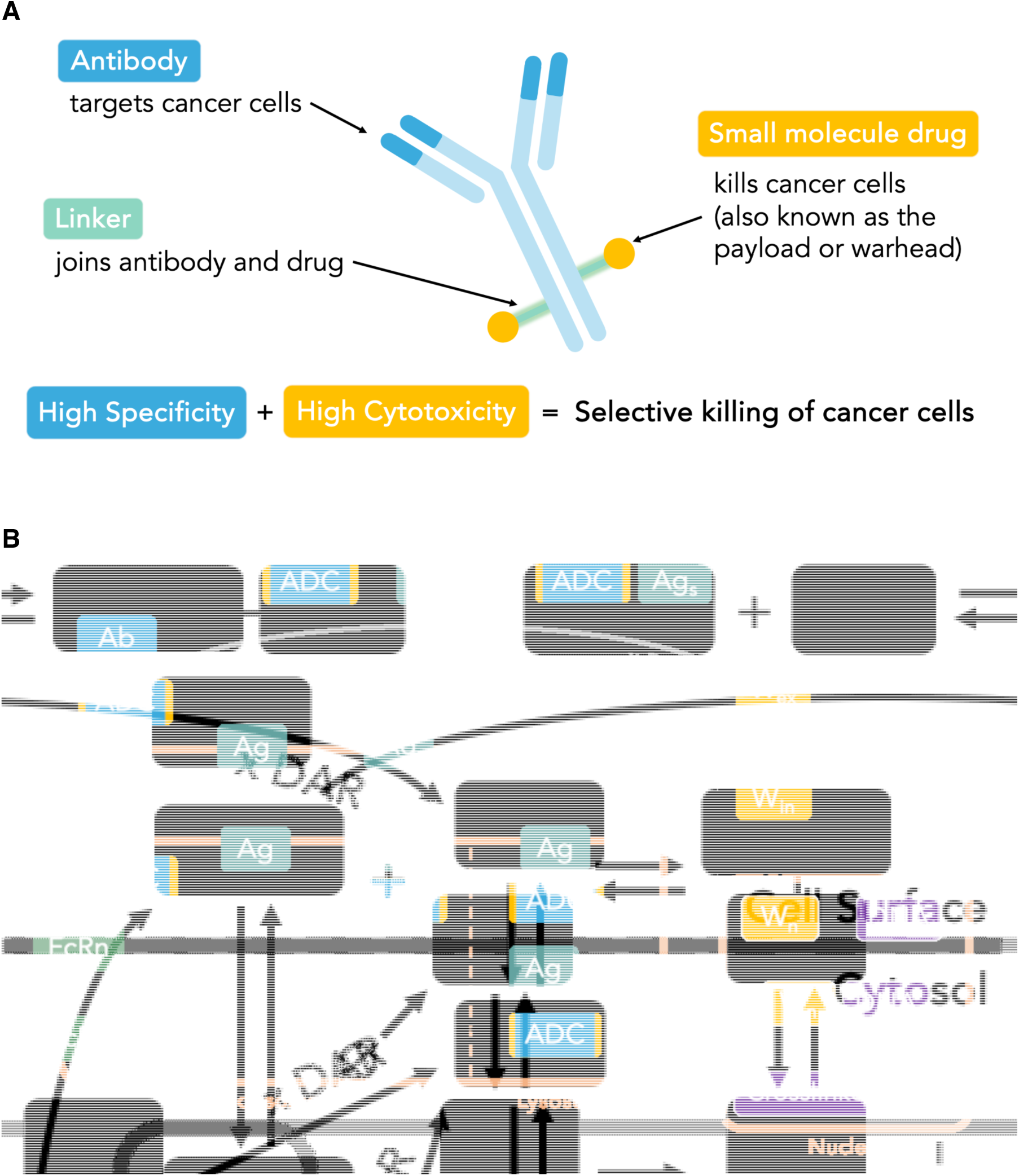
ADC structure and schematic of model mechanisms. **(A)** Simplified structure of an antibody-drug conjugate (ADC), showing the three key components of the molecule. The colors of each component correspond to those used in the model schematics (Figure 1B and elsewhere). (**B)** Computational model schematic showing the network of reactions. The antibody component of the ADC binds to the tumor-specific antigen (Ag) expressed on the cell surface where it is accessible to the extracellular ADC. Ag-bound ADC can be internalized and recycled, and both intracellular and extracellular ADC can release warhead (W) in the model as shown. Intracellular warhead can enter the nucleus (its site of action) or can leave the cell. We developed a mechanistic ordinary differential equation model based on this schematic that tracks the concentrations of each of the key components shown (filled boxes) and how those concentrations are impacted by the rates of the processes (arrows). The complete set of equations is given in **Supplemental Methods S1**, and the parameters are described in **Tables 1-5**. A more detailed schematic with processes labeled can be found in **Figure S1**. DAR = drug-antibody ratio.

As a specifically designed, targeted drug delivery vehicle to selected cells, ADCs have revolutionized the treatment of certain cancers, with 13 ADCs approved by the FDA for use in oncology [23–25]. With over half of these approvals occurring within the last 5 years, interest in ADCs remains high, with over 160 candidates currently in clinical trials [26]. In this study, we focus on a computational systems pharmacology model of an ADC (MEDI2228) developed by AstraZeneca. MEDI2228 contains two molecules of a pyrrolobenzodiazepine (PBD) dimer warhead (SG3199), covalently linked via site-specific conjugation at the cysteine residues to an anti-BCMA IgG1 monoclonal antibody via a protease-cleavable valine-alanine dipeptide linker [27]. PBD dimers act by covalently binding to the minor groove of DNA to induce DNA crosslinking, thereby preventing cell division and ultimately causing apoptosis [28,29]. MEDI2228 targets B-Cell Maturation Antigen (BCMA) [27,30], which is expressed on B-cells and hematological cancers such as multiple myeloma [31].

In order to improve our understanding of this ADC and ultimately enable improved ADC development, we have constructed a QSP model parameterized using optimization to *in vitro* data, exploring model parameters via sensitivity analysis and developing insights based on model simulations. This model can serve as a platform for studying different ADCs; here, we use the MEDI2228 as the representative ADC. Previous computational models of ADCs have included different levels of detail of cellular mechanisms [32]. Some earlier mathematical models of ADCs were semi-mechanistic at the *in vitro* scale and focused on a limited number of key processes, including ADC-antigen (Ag) binding, internalization, and warhead efflux ([33–35]; reviewed in [32]). Later ADC models became more complex with more detailed mechanisms such as trafficking, degradation, and receptor recycling ([36–38]; reviewed in [32]). Here, we incorporate key elements from those models, but also expand upon them. There have not yet been published computational models for PBD ADCs, and so we add mechanistic detail specific to PBD ADCs (such as warhead-DNA binding and DNA crosslinking), as well as FcRn recycling and binding to both soluble and membrane-associated antigen in order to capture a broader scope of the biology, particularly for endogenous receptors as antigens. Notably, we built in tracking of various extracellular and intracellular sources of warhead into the model to better visualize the movement of warhead at the subcellular and multicellular levels. Using MEDI2228 as the representative ADC, this computational model allows us to capture specific insights while containing generalizable and modular components that can be modified to describe other ADCs.

## Methods

### Computational model construction

We have constructed a QSP-type model composed of a set of coupled, deterministic, nonlinear, ordinary differential equations (ODEs), each describing the rate of change over time in concentration of a molecule or molecular complex in the system (see schematic, **Figure 1B**). The model includes multiple compartments representing the extracellular space (including the cell surface), endosomes and lysosomes, cytosol, and the nucleus, along with a clearance compartment to track degraded antigen and antibody. The concentrations of molecules in each compartment are assumed to be well-mixed. The same molecule in different locations (e.g. extracellular warhead, intracellular warhead, nuclear warhead) each has its own equation, and these equations are coupled through the terms that describe the processes of transport or interaction. Model equations are included in the supplemental information (**Supplemental Methods S1**) and are in units of nM×hr^-1^ except for the equations for the concentrations of cell surface molecules: antigen (Ag); ADC-bound antigen (ADC-Ag); and antibody-bound antigen (Ab-Ag), which are in nM×cell^-1^×hr^-1^ and represent the concentration at the single cell level (described in the next section; this is done to enable the equations to deal with the proliferation and death of cells). Some of the processes included here have been studied and modeled by others for different ADCs [32,33,36,37,39]. These key biological processes are described in the following sections, and their corresponding model parameters were obtained from multiple sources, as appropriate (**Table 1**). The model allows for dosing of various therapeutic modalities, including standard ADC, isotype control ADC (which cannot bind to the target Ag), unconjugated Ab (i.e. no warhead), and free warhead (i.e. no Ab).

**Table 1.** Volume parameter values for the computational model. Because the model includes multiple different compartments - extracellular media, cytoplasm, endosome, nucleus - we include here the volumes of each and the sources of the values.

| Parameter | Description | Value | Unit | Source |
| --- | --- | --- | --- | --- |
| **Volumes** |  |  |  |  |
| $V_{\text{cell}}$ | Volume of a single cell (assume to be cytoplasm) | $1.00\text{E-}12$ | L/cell | [58] |
| Nucleus Percent OfCell | Percentage of cell made up of nucleus | 10 | % | [59] |
| $V_{\text{nucleus}}$ | Volume of nucleus in a single cell | $1.00\text{E-}13$ | L/cell | Calculated |
| Endosomes Percent OfCell | Percentage of cell made up of endosomes and lysosomes | 5 | % | [60,61] |
| $V_{\text{endo}}$ | Volume of endosomes in a single cell | $5.00\text{E-}14$ | L/cell | Calculated |
| $V_{\text{media}}$ | Volume of extracellular media | $100\text{E-}06$ | L | Varies based on experimental conditions |

**Table 2.** Initial conditions for the computational model. The parameter values used to simulate the experimental conditions. Parameters for the anti-BCMA PBD ADC and its related experiments are listed here, including sources of the values (calculated, optimized, estimated, or obtained from the literature).

| Parameter | Description | Value | Unit | Source |
| --- | --- | --- | --- | --- |
| **Experimental Conditions: MEDI2228 Cytotoxicity Assay** |  |  |  |  |
| PBD0 | Initial concentration of PBD warhead | 7.8E-08 - 1E+02 | nM | Varies based on experimental conditions (see footnote A) |
| ADC0 | Initial concentration of ADC | 1.86E-05 - 3.30 | nM | Varies based on experimental conditions (see footnote B) |
| Ag0 | Number BCMA receptors per cell for NCI-H929 | 3.145E-11 | nmol/cell | Calculated using 18,931 receptors/cell [27] (see footnote C) |
| DNA0 | Initial amount of DNA binding sites | 2.49E-09 | nmol | Estimated (see footnote D) |
| ADC_MW | MEDI2228 molecular weight | 151,614 | Da (g/mol) | [27] |
| BCMA_MW | BCMA molecular weight | 5,300 | Da (g/mol) | [62] |
| FcRn0 | Initial amount of FcRn | 2.49E-09 | nmol | [42] |
| Ag_s0 | Soluble BCMA receptors | 0, 7.5, 28, 72 10<sup>-5</sup> | g/L | [27] |
| q | Infusion rate of ADC | 0 | nmol/hr | No infusion |
| Ncell | Initial number of cells | 5,000 - 10,000 | cells | Varies based on experimental conditions; calculated from cell density |

$N_A = 6.02 \times 10^{23}$ molecules/mol (Avogadro's number)
- A. Initial PBD concentrations include: 7.81E-08, 1.56E-06, 3.12E-05, 6.25E-04, 0.0125, 0.25, 5, 100 nM - B. Initial ADC concentrations include: 1.86E-05, 5.59E-05 1.68E-04 5.03E-04 1.51E-03 4.52E-03, 0.014, 0.041, 0.12, 0.37, 1.10, 3.30 nM - C. $Ag_0 = (\# \text{ receptors/cell}) / N_A \times 10^9$ - D. $DNA_0 = (3E+09 \text{ base pairs of DNA}) / (10 \text{ base pairs per turn}) \times (50\% \text{ of grooves created by each turn are expected to be minor grooves}) \times (1\% \text{ of genome consisting of coding regions of DNA}) / N_A \times 10^9$

**Table 3.** Parameter values for the mechanistic computational model, part 1 - antibody-antigen binding. Parameters for the anti-BCMA PBD ADC are listed here, including sources of the values (calculated, optimized, estimated, or obtained from the literature).

| Parameter | Description | Value | Unit | Source |
| --- | --- | --- | --- | --- |
| **ADC and Ab Binding/Unbinding to Antigen** |  |  |  |  |
| $k_{onAg}$ | Binding to human membrane-bound BCMA | 1.34 | $nM^{-1} hr^{-1}$ | Assumed to be the same as $k_{onAg\_s}$ |
| $k_{offAg}$ | Unbinding from human membrane-bound BCMA | 4.16268 | $hr^{-1}$ | Calculated using $K_D = 3.1$ from Supplementary Table S4 in [27] |
| $k_{onAg\_s}$ | Binding to human monomeric sBCMA | 1.34 | $nM^{-1} hr^{-1}$ | Unpublished data from [27], measured via surface plasmon resonance |
| $k_{offAg\_s}$ | Unbinding from human monomeric sBCMA | 81.72 | $hr^{-1}$ | Unpublished data from [27], measured via surface plasmon resonance |

**Table 4.** Parameter values for the mechanistic computational model, part 2 - trafficking. Parameters for the anti-BCMA PBD ADC are listed here, including sources of the values (calculated, optimized, estimated, or obtained from the literature).

| Parameter | Description | Value | Unit | Source |
| --- | --- | --- | --- | --- |
| **Internalization/Recycling/Trafficking** |  |  |  |  |
| **Of ADC-Ag Complex** |  |  |  |  |
| $k_i$ | ADC-Ag complex internalization rate | 0.09437 | hr<sup>-1</sup> | Optimized |
| $k_e$ | ADC-Ag complex Exit rate from endosome | 2.961 | hr<sup>-1</sup> | Optimized |
| $f_{rec}$ | Endosomal recycling fraction for ADC-Ag complex | 0.4344 | Unitless | Optimized |
| **Of Antigen** |  |  |  |  |
| $k_{iAg}$ | Ag internalization rate | 0.09437 | hr<sup>-1</sup> | Fixed to $k_i$ |
| $k_{eAg}$ | Ag exit rate from endosome | 2.961 | hr<sup>-1</sup> | Fixed to $k_e$ |
| $f_{recAg}$ | Endosomal recycling fraction for antigen | 0.4344 | Unitless | Fixed to $f_{rec}$ |
| **Of ADC** |  |  |  |  |
| $k_{iADC}$ | ADC internalization rate (Pinocytosis) | 1.2095 | hr<sup>-1</sup> | Optimized |
| $k_{eADC}$ | ADC Exit rate from endosome | 25 | hr<sup>-1</sup> | [42] |
| $k_{recADC}$ | Endosomal recycling rate for ADC | 9.2597 | hr<sup>-1</sup> | Optimized |
| **FcRn Binding** |  |  |  |  |
| $k_{onFcRn}$ | ADC binding to FcRn | 0.1261 | nM<sup>-1</sup> hr<sup>-1</sup> | Calculated using $k_{offFcRn}$ and $K_D = 648$ nM (unpublished data from [27,30]) |
| $k_{offFcRn}$ | ADC unbinding from FcRn | 81.72 | hr<sup>-1</sup> | Assumed to be the same as unbinding rate from target receptor |
| **Degradation** |  |  |  |  |
| $k_{deconjugation}$ | Extracellular ADC Deconjugation (cell culture media / mouse serum) | 0 or 3.2617 $\times 10^{-4}$ | hr<sup>-1</sup> | Assumed *in vitro* stability of ADC in media / Optimized |
| **Antigen Production** |  |  |  |  |
| $k_{AgProduction}$ | Antigen production rate | 1.678e-12 | nmol cell<sup>-1</sup> hr<sup>-1</sup> | Optimized |

**Table 5.** Parameter values for the mechanistic computational model, part 3 - warhead and tumor cell dynamics. Parameters for the anti-BCMA PBD ADC are listed here, including sources of the values (calculated, optimized, estimated, or obtained from the literature).

| Parameter | Description | Value | Unit | Source |
| --- | --- | --- | --- | --- |
| **Warhead release** |  |  |  |  |
| DAR | Drug-to-Antibody Ratio | 2 | Number per Ab | Known value for MEDI2228 |
| **Warhead Movement** |  |  |  |  |
| $k_{\text{efflux}}$ | Efflux | 0.1384 | hr<sup>-1</sup> | Calculated based on PBD half-life from [41] |
| $k_{\text{influx}}$ | Influx | 11.55 | hr<sup>-1</sup> | Calculated based on PBD half-life from [41] |
| $k_{\text{in\_nuc}}$ | Trafficking to Nucleus | 0.1384 | hr<sup>-1</sup> | Calculated based on PBD half-life from [41] |
| $k_{\text{out\_nuc}}$ | Trafficking out of Nucleus | 11.55 | hr<sup>-1</sup> | Calculated based on PBD half-life from [41] |
| **PBD Mechanism of Action** |  |  |  |  |
| $k_{\text{onDNA}}$ | Binding to DNA | 0.06 | nM<sup>-1</sup> hr<sup>-1</sup> | Estimated from reaction rate constant range for anthramycin binding |
| $k_{\text{offDNA}}$ | Unbinding from DNA | 0.006 | hr<sup>-1</sup> | Calculated from dissociation constant and $k_{\text{onDNA}}$ |
| $K_D$ | Dissociation constant of PBD dimer binding DNA | 0.1 | nM | Estimated from $K_D$ range for PBDs |
| $n$ | Hill Coefficient for DNA Crosslinking | 1 | Unitless | Fixed at 1 |
| Crosslink IC50 | PBD dose producing 50% DNA crosslink formation | 1.24 | nM | Optimized |
| $K_A$ | Concentration of Wn-DNA at PBD dose producing 50% DNA crosslink formation | 121.6738 | nM | Calculated based on Crosslink IC50 |
| $k_{\text{kill}}$ | Cell killing | 0.11708 | hr<sup>-1</sup> | Optimized |
| **Tumor Cell Characteristics** |  |  |  |  |
| $t_{1/2}$ | NCI-H929 cell doubling time | 50 | hr | [63] |
| $k_g$ | Net growth rate of cancer cell line | 0.0139 | hr<sup>-1</sup> | Calculated from NCI-H929 doubling time; $k_g = \ln(2) / t_{1/2}$ |

### Model equations describe concentrations at the single cell level while reflecting the changing cell population

In order to take the effects of a dynamic cell population into account (i.e. to account for the changing cell density due to proliferation and killing) while maintaining model flexibility, we crafted the equations for molecules on the cell surface (Ag, ADC-Ag, and Ab-Ag) at the single cell scale with units of nM×cell^-1^×hr^-1^, enabling easy conversion between the single cell-level and population-level concentrations simply by multiplying by cell population. Equations describing ADC, Ab, soluble BCMA (Ag_s_), and extracellular warhead (W_ex_) are in units of nM×hr^-1^ relative to the media (extracellular volume) of the cell culture system. Within these equations describing concentrations in the extracellular volume (notably ADC, Ab, and W_ex_), any processes involving interactions with the cells (binding, unbinding, pinocytosis, FcRn recycling, influx, and efflux) are scaled by the cell population to ensure the reactions and effects across all cells are captured. For example, for ADC binding to Ag, more cells means more Ag, and therefore more depletion of ADC from the media. The cells equation has the units of (# of cells)*hr^-1^, and this dynamic number of surviving cells is incorporated into the model equations where processes need to be scaled by the cell population. All remaining equations describing entities in the endosomal/lysosomal space, cytoplasm, or nucleus are in units of nM×hr^-1^ relative to the volume of their respective subcellular locations. These intracellular concentrations remain the same, regardless of examining the concentration within a single cell or across all cells (due to the proportional increase in volume per cell). The implicit assumption here is that when cells die, the molecules in those cells are lost, and thus the net effect on the tracked intracellular concentration in the remaining cells is zero. By building the equations in this way, we can track the effects of both cell proliferation and cell killing, which is important in comparing simulations to experiments with and without treatment (as cells are both proliferating and being killed).

### Antigen Binding and Production

Binding of the ADC to its target antigen (Ag) is the first step to efficacy; the binding affinity of the antibody affects the ADC’s ability to target receptors on tumor cells and normal cells that have different levels of expression of the receptors. Here, rate constants for binding (k_onAg_) and unbinding (k_offAg_) between the ADC and Ag on the cell surface were derived from known values for the binding rate constant to monomeric BCMA (AstraZeneca, previously unpublished data generated for ref [27]) and the dissociation constant of MEDI2228 to membrane-bound BCMA [27]. We assume unconjugated antibody (Ab) binds to and unbinds from Ag with the same rate constants, as conjugation of the warhead occurs in the constant region of the Ab, rather than in the variable region that is involved in Ag binding. We also included binding (k_onAgs_) and unbinding (k_offAgs_) of the ADC/Ab to soluble BCMA (Ag_s_) in the extracellular space to explore *in vitro* effects of soluble BCMA as a potential sink for anti-BCMA ADCs, using measured values for binding and unbinding rate constants of MEDI2228 to soluble (monomeric) BCMA (AstraZeneca, previously unpublished data generated for ref [27]). For simulations involving the isotype control ADC, the rate constants for binding (k_onAg_) and unbinding (k_offAg_) were set to zero because it does not bind the target antigen. The rate of antigen production (k_AgProduction_) was represented as a zeroth order process, adding Ag to the cell surface at a constant rate in nmol×cell^-1^×hr^-1^; the value of k_AgProduction_ was optimized to maintain the initial antigen concentration in the system in the absence of any drug (but with internalization, trafficking, recycling, and degradation processes functioning) for a static cell population (k_g_ = 0) after 7 days. A pre-simulation that calculates the initial Ag_endo_ and Ag_lys_ concentrations is also built into the initialization of the model by simulating the static cell population without drug up to 7 days to represent the steady state, and taking the respective final concentrations as the initial values for further simulations.

### Trafficking of ADC/Ab: Internalization, Recycling, Degradation, and Warhead Release

ADC uptake into cells is key to efficacy, as internalization enables the cytotoxic payload to be released and reach the cell nucleus. We added ADC and Ab internalization to the model as first-order processes, including both receptor-mediated endocytosis (k_i_) and non-receptor-mediated pinocytosis (k_iADC_). Including pinocytosis enables the simulations to account for weak cell killing by isotype control ADCs.

Following receptor-mediated internalization, the ADC-Ag complex may be recycled back to the cell surface or trafficked from the endosome to the lysosome; these processes are represented using a fraction recycling parameter (f_rec_) and an endosomal exit rate constant (k_e_). While the endosomal and lysosomal spaces are treated as the same compartment within the model, the molecules in the lysosome are represented with separate equations, and degradation of molecules in the lysosome is assumed to proceed irreversibly. Unbound Ag can similarly undergo internalization, recycling, and lysosomal trafficking (its loss to degradation generally being balanced by the constant production of new Ag). Parameters for the receptor-mediated endocytosis pathway (k_iAg_, f_recAg_, k_eAg_) were optimized to and validated against experimental data measuring the *in vitro* cytotoxicity for MEDI2228 [30].

Conversely, parameters for the non-receptor-mediated pathway involving unbound ADC (k_iADC_, k_recADC_, k_eADC_) were fit using *in vitro* cytotoxicity data for isotype control ADCs that can only enter the cell via nonspecific uptake or pinocytosis (AstraZeneca, previously unpublished data generated for ref [30]). ADC pinocytosis occurs when cells sample the surrounding fluid, resulting in an exchange of volumes between the extracellular space near the cell and the intracellular endosomes. However, in an *in vitro* system, the volume of media in the extracellular space is much greater than the volume of the intracellular space. As a result, when the extracellular ADC concentration is multiplied by a volume correction term to represent the transition from the extracellular to the endosomal compartment, this can lead to an overrepresentation of the amount of ADC uptake via pinocytosis in the model. Thus, in this model, we multiply the pinocytosis rate constant by the difference in extracellular and endosomal ADC concentrations, i.e. [ADC_ex_ - ADC_endo_], such that the exchange of volumes is consistent and the concentration of the ADC inside and outside the cell will move towards an equilibrium.

Unbound ADC that is internalized via pinocytosis is also subject to recycling and lysosomal trafficking but must first bind to the neonatal Fc receptor (FcRn) to be recycled. Rate constants for ADC binding (k_onFcRn_) and unbinding (k_offFcRn_) to FcRn were measured directly using surface plasmon resonance (AstraZeneca, previously unpublished data generated for [27]).

Following trafficking to lysosomes, both the ADC-Ag complex and unbound ADC in the model are degraded, and the PBD warhead is released into the cytosol of the cell in accordance with its DAR value (2:1 ratio of warhead to ADC for MEDI2228). Degradation rates may correspond with the amount of warhead available at the site of action, and thus impact efficacy.

### Efflux, Influx, and Warhead Trafficking

After release into the cytosol (following lysosomal degradation of the ADC), there are multiple potential paths for the warhead. Efflux and influx of the warhead reflect its propensity to stay within the cell or to escape the cell and cause bystander killing [40,41]. Similarly, the warhead can diffuse to and from the site of action in the nucleus of the cell. For this model, we assumed that these processes only include passive diffusion, and the parameters for efflux and influx (movement out of and into the cell) were calculated based on estimated PBD half-life values from computational analyses based on the membrane permeability and partition coefficient [41]; the values are in a similar range to those for other warheads [41]. The parameter values for influx and efflux are based on the partition coefficient of PBD between intracellular and extracellular spaces, which assumes similar volumes across the membrane between the partitions. However, in the *in vitro* system, the volumes between the extracellular space and intracellular cytosol differ by several orders of magnitude, and the volume correction term for movement of warhead from the extracellular media to the cytoplasm (V_media_/V_cytoplasm_) amplifies the resulting cytosolic concentration. Thus, we introduced an additional volume adjustment term (V_cytoplasm_/V_media_) to account for these volume differences, which cancels with the original volume correction term for influx. This helps to maintain the partition coefficient while also better representing the warhead locally available near the cell surface (rather than warhead in the media that is too far away to enter the cell), and notably maintains the expected behavior of warhead partition at the lower extracellular volumes typical of tissues *in vivo*.

### Warhead Binding and DNA Crosslinking

Once the warhead reaches the nucleus (the site of action for PBDs), it can bind to minor groove binding sites on DNA. Warhead-DNA (W_n_-DNA) binding and unbinding rate constants were estimated based on the picomolar-level affinity typically observed for PBD warheads. This binding step induces the formation of DNA crosslinks; we used the Hill equation to represent the fraction of crosslinked DNA as a function of the level of warhead in the nucleus bound to DNA.

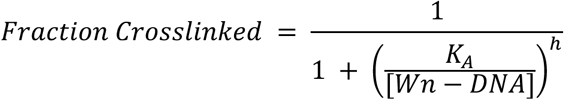

Values for the Hill coefficient (h) and the W_n_-DNA concentration producing 50% crosslinking (K_A_) were optimized to experimental data measuring dose-response DNA interstrand crosslink formation as a result of PBD binding in NCI-N87 cells [29] (described in *Results*).

### Cell Growth and Inhibition

The *in vitro* tumor cell population in the model proliferates at the net growth rate (k_g_), based on the known doubling times for the specific cell line (NCI-H929) in the experiment being simulated. Cell killing is dependent on crosslink formation and uses rate constants obtained from optimization to data from dose-response *in vitro* cytotoxicity studies in the NCI-H929 cell line (AstraZeneca, previously unpublished data). This parameter is an indicator of warhead potency and can be used to explore the effect of potency on both efficacy and toxicity (via bystander effect) with the cell population. Combining both cell growth and killing, our model equation for cells begins at the initial number of cells and calculates change in the cell population over time. This enables conversion from single cell concentrations to population-level effects across the entire cell culture system, as processes involving cell surface interactions (binding, unbinding, pinocytosis, FcRn recycling, influx, and efflux) are scaled by cell population, increasing and decreasing alongside the number of cells.

### Simulation

Using MATLAB to encode the model equations, we used ODE solver ode23s to run individual numerical simulations for various *in vitro* cell culture experiments in the NCI-H929 cell line by replicating experimental conditions *in silico*. For instance, known BCMA receptor density values for NCI-H929 cells (malignant B lymphocytes) were used in simulations of the anti-BCMA PBD ADC (AstraZeneca, previously unpublished data generated for ref [30]). Simulation times and initial conditions, such as ADC levels in the culture media, were specific to the experiment being simulated and the values used in the simulations were the same as those in the experiments. Parameter values for each model were measured from experimental data, pulled from literature, estimated, or optimized as described below (see *Results* and **Table 1**).

### Optimization

For parameters that were not measured or could not be found through literature, we used the nonlinear least squares optimization function (*lsqnonlin*) in MATLAB to identify optimal values by comparing simulation results to corresponding experimental data and minimizing the squared norm of the residual (i.e. the difference between simulation predictions and experimental results), as described in the Results section (these parameters are labeled as “Optimized” in Tables 3-5). Notably, we used a sequential parameter optimization process that takes advantage of multiple types of experimental data, some previously published and some provided by AstraZeneca. The data from AstraZeneca measured *in vitro* cytotoxicity (cell survival percentages) at different doses of (1) the PBD warhead (SG3199), (2) an isotype control ADC that also has 2 PBD dimers but cannot bind to BCMA, and (3) MEDI2228, the standard PBD ADC that binds to BCMA. We performed each optimization step with one hundred initial guesses for each parameter over a designated parameter value range, randomly generated using the Latin Hypercube Sampling function (*lhsdesign*) in MATLAB.

### Sensitivity Analysis

We used sensitivity analyses to examine the impact of the parameters on the system outputs. For the univariate local sensitivity analyses, each parameter value was increased (one at a time) by 10%, and the simulations were run under the same conditions (time = 96 hours, ADC dose = 4.1 x 10^-2^ nM, and 1.9 x 10^4^ receptors per cell) for each varied parameter to determine the sensitivity, i.e. the effect on key outputs. Here, sensitivity is defined as the percent change in area under the curve (AUC) of the output normalized to the percent change in parameter value (10%). The AUC was calculated by integrating the concentration of key outputs (e.g. warhead, crosslinking, or cell population) over time. Parameters with values of zero (i.e. q and k_deconjugation_) were excluded from the analysis. Similarly, we conducted univariate local sensitivity analyses at different time points following ADC addition, different antigen expression levels, and different ADC doses to explore how the parameter sensitivities behaved under different conditions. Finally, we performed a univariate global sensitivity analysis by varying the final identified parameter values over a range from two orders of magnitude below to two orders above the original values. Parameters with values of zero (i.e. q and k_deconjugation_) or with values that could not be reasonably varied over multiple orders of magnitude (e.g. volumes and fractions) were excluded from the analysis. We visualized these perturbations by calculating the root sum of squares error (using the MATLAB function *rssq*) between the model predictions and the experimental *in vitro* cytotoxicity data (both measured at 96 hours following MEDI2228 dosing).

## Results

### Sequential parameterization of the model by modular simulation

We have constructed a detailed and modular computational model of MEDI2228 and its associated biological mechanisms. Some parameters for the model were derived from literature (including from previous models of ADCs and antibodies generally) or directly from experimental measurements (**Table 1**), while other parameters, described here, were obtained by optimization of simulation results to experimental data.

During the model parameterization and optimization process, we first sought to identify the warhead-dependent parameters to match *in vitro* experimental data using the SG3199 warhead, as these are furthest downstream mechanisms and have the fewest dependent variables. Moreover, the warhead is the entity that ultimately drives the cytotoxic effect of the ADC. Once the warhead-dependent parameters were identified, we moved on to optimize ADC/Ab-dependent parameters which govern the upstream mechanisms tied to the Ab portion of the ADC, calibrating to *in vitro* experimental data that used the entire ADC, first using an isotype control ADC and then the specific anti-BCMA ADC. These newly-identified model parameters are described in the following sections.

### DNA crosslinking represented using the Hill Equation

Due to the limited data on PBD-DNA binding and unbinding kinetics, we sought alternative options to parameterize the portion of the model involving pharmacokinetics and pharmacodynamics (PKPD) of the warhead (**Figure 2A**). To optimize parameters for the Hill equation used to represent the fraction of crosslinked DNA, we used experimental data on DNA interstrand crosslink formation over a range of PBD doses [29]. Though the cell line used for these experiments (NCI-N87 cells) differed from the NCI-H929 cells that are the focus of the other simulations, it provided the best option given the data available. We explored multiple possible combinations of values of the two key parameters, the Hill coefficient (n) and the concentration of Wn-DNA at the PBD dose producing 50% crosslinking (K_A_), compared the predictions to the experimental data for crosslinking level, and calculated the sum of square error over the range of possible values for n and K_A_ (**Figure S2A**). Though the goal is typically to obtain the combination of values that produces the smallest error, we noted that the difference in error across these values is relatively small, particularly in varying n. Therefore, we fixed the Hill coefficient to 1 to obtain a simpler, more parsimonious model while still maintaining a good fit to the experimental data. Simulations of dose response experiments at various values of K_A_ demonstrate that fixing n to 1 provides a reasonable fit of the experimental data (**Figure S2B**). After optimizing for K_A_ at n = 1 (**Figure 2B**), we validated the chosen crosslinking parameters by simulating a time course experiment and comparing the results to the experimental data (**Figure 2C**).

**Figure 2.**
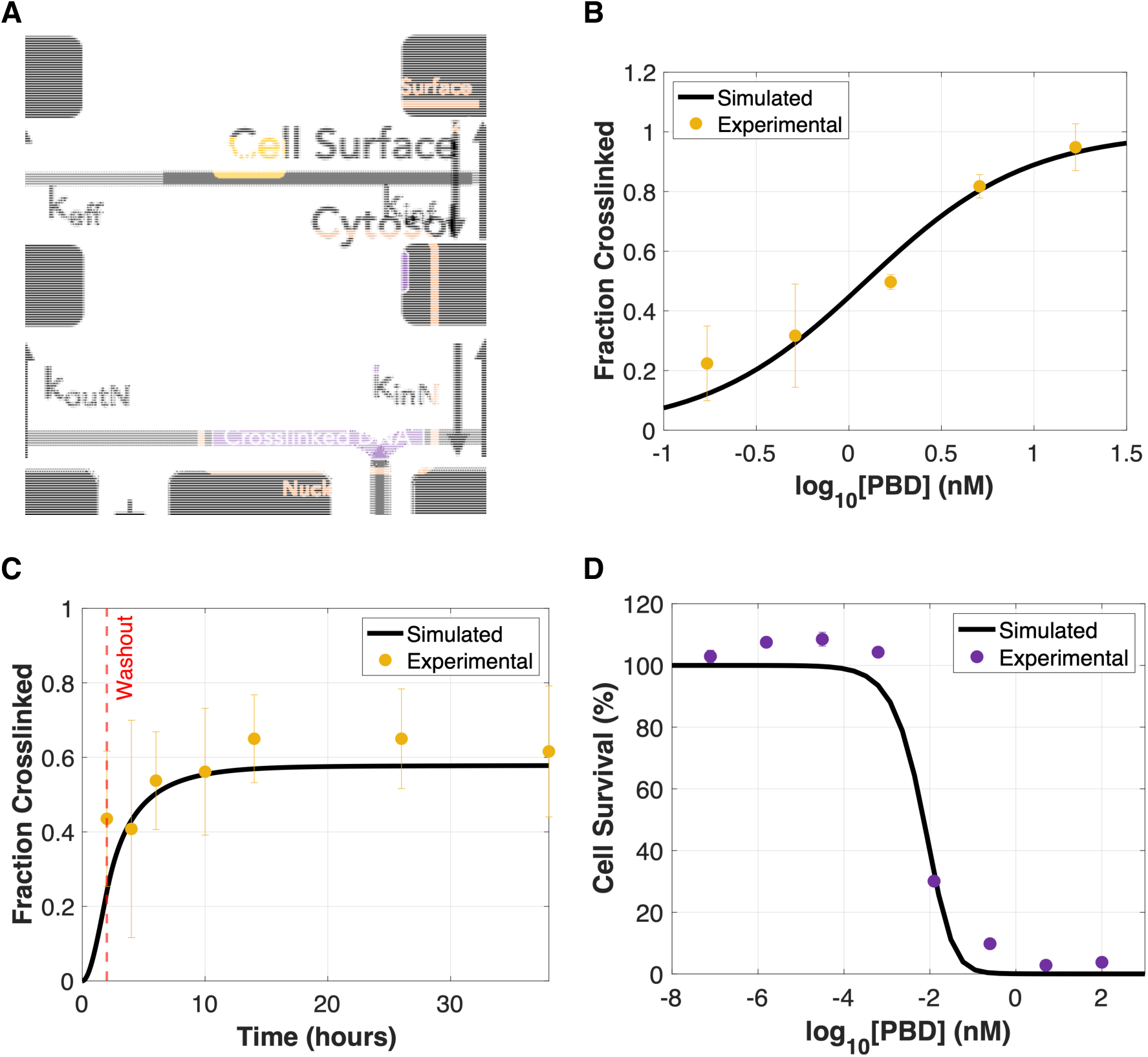
Warhead module and identification of warhead-dependent parameters. **A**, Schematic showing the module of PBD dimer warhead pharmacokinetics and pharmacodynamics within the larger computational model framework. We can simulate isolated modules of the larger model to facilitate comparison to experiments that only involve the components in those modules. This modular assembly and parameterization of the model facilitates use of multiple different data sets. **B,** Comparison of simulation results (line) and experimental data (dots) for DNA crosslink formation in response to a 2 hour treatment with SG3199 (the PBD dimer warhead used in MEDI2228) followed by incubation for 24 hours in drug-free media. DNA crosslink formation in the model is represented with a Hill term, using optimized values for *KA* and *n*. **C,** Validation of module parameters by comparing time course simulation results (line) to experimental data (dots) of DNA crosslink formation following a 2 hour treatment with SG3199, then an incubation to 36 hours in drug-free media. The washout time is indicated by the vertical red dotted line. **D,** Optimization of cell killing rate (kkill) using experimental data (dots) from an NCI-H929 cell line results in a reasonable fit of simulation-predicted cell survival (line) at varying SG3199 doses. The error bars for the experimental data in panel D are very small and not visible for most of the data points.

### Cytotoxic effect of warhead optimized to cell survival data

Following the identification of the DNA crosslinking parameters, the rate of cell killing, as a function of crosslink formation, was optimized to experimental data measuring the *in vitro* cytotoxicity in NCI-H929 cells at varying doses of the SG3199 warhead (**Figure 2D**). Using warhead-specific experimental data enables the isolation of the cytotoxic effect of the PBD warhead, as opposed to the entire ADC.

### Non-receptor-mediated trafficking parameters identified using isotype control ADC cytotoxicity data

As the isotype control ADC does not bind to the target receptor, we can assume that any uptake of isotype control ADC into the cell is from non-receptor-mediated internalization pathways, such as pinocytosis. Thus, we were able to use the model simulations to identify the parameter values for the non-receptor-mediated internalization (pinocytosis) rate constant, endosomal exit rate constant, and FcRn recycling rate constant for the ADC by optimizing to data for *in vitro* cytotoxicity in NCI-H929 cells across different isotype control ADC doses at 9, 24, 48, and 72 hours (**Figure 3A**). Though we initially fit all 3 parameters simultaneously, we found that the parameter values were not uniquely constrained and several parameter sets were possible (**Figure S3**). Thus, we fixed the endosomal exit rate constant based on a literature value [42], while optimizing the remaining parameter values for pinocytosis and FcRn recycling rate constants, resulting in a better-constrained parameter space (**Figure S4**). Of the parameter values with the highest frequency, we selected the parameter set that gave the lowest cost (sum of squared residuals). Validation to a separate *in vitro* experiment in the same cell line, which also measured cell survival over varying doses of isotype control ADC but at a later time point (96 hours), shows that these parameters are able to reflect the experimental data accurately (**Figure 3B**).

**Figure 3.**
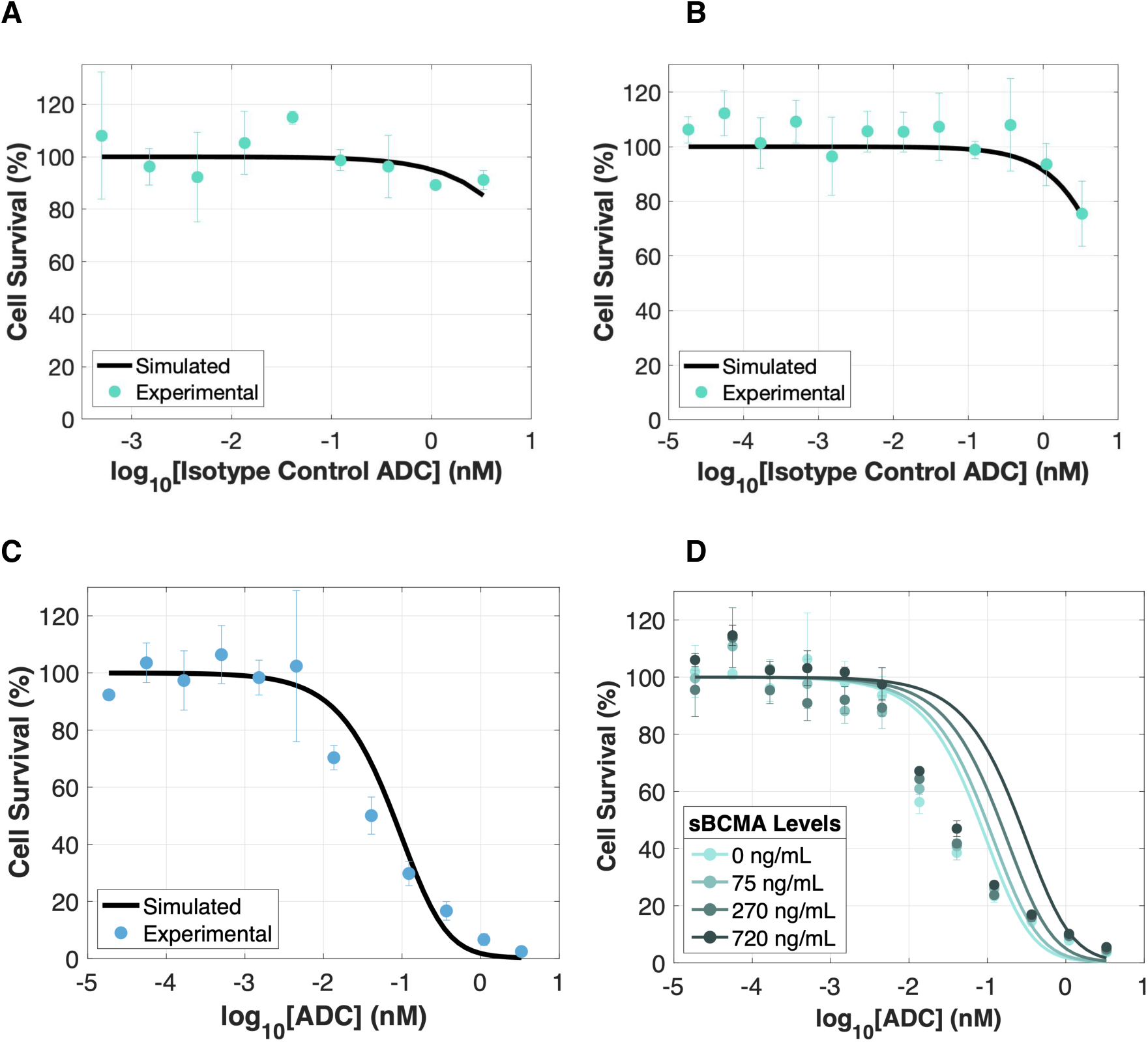
Optimization and validation of full model to *in vitro* cytotoxicity experimental data at varying doses of isotype-control ADC and antigen-targeting ADC. **A,** Parameters for non-specific uptake rates were identified by fitting to *in vitro* cytotoxicity data (dots) in NCI-H929 cells at 72 hours after exposure to varying doses of isotype control ADC; since the isotype control does not bind to the antigen, cell death would be due to non-specific uptake. The line shows the predicted cell survival following optimization of parameters for non-receptor-mediated ADC internalization (*ki,ADC*), endosomal exit rate (*ke,ADC*), and recycling (*krec,ADC*). **B,** Validation by comparing separate experimental data (dots) in the same cell line to simulation results (line) of cell survival at 96 hours after isotype control ADC exposure. **C,** Optimization of receptor-mediated endocytosis and trafficking parameters—including internalization of Ag and ADC-Ag (*ki*, *ki,Ag*), endosomal exit rate constants (*ke*, *ke,Ag*), and the fraction of recycling (*frec*, *frec,Ag*), as well as antigen production (*kAgProduction*)—to *in vitro* cytotoxicity data (dots) for NCI-H929 cells at 96 hours following administration of MEDI2228. **D,** Validation by comparing experimental data (dots) to simulation results (lines) for soluble antigen confirms that the addition of different levels (0, 75, 270, and 720 ng/mL) of soluble BCMA (sBCMA) does not strongly impact MEDI2228 efficacy, and that . the simulations reflect the slight dose-dependence trends of the experimental data. Both the simulations and experiments show that while the cell survival curves shifted slightly to the right (indicating a slight increase in IC50s) at higher levels of sBCMA, the overall effect of soluble receptors has minimal impact on the ADC’s efficacy.

### Receptor-mediated internalization pathway and receptor trafficking parameters fit using standard ADC cytotoxicity data

Finally, the remaining parameters describing the internalization, endosomal exit, and recycling of both the ADC-Ag complex and unbound Ag were fit to *in vitro* cytotoxicity data in NCI-H929 cells at varying doses of the anti-BCMA PBD ADC at 9, 24, 48, 72, and 96 hours (**Figure 3C**). We nested the optimization of the rate of antigen production within the outer optimization function to ensure steady state levels typical of observed antigen levels were reached in the absence of ADC. After initially optimizing for all 6 trafficking parameters simultaneously, we found that the parameter space was not well-constrained (**Figure S5**). In addition, inconsistent differences between the rate constants for Ag and ADC-Ag complex were observed, and thus we linked the parameters for ADC-Ag complex and unbound Ag together (i.e. k_i_ = k_iAg_, k_e_ = k_eAg_, and f_rec_ = f_recAg_), simplifying the optimization to 3 parameters and resulting in a narrower parameter space (**Figure S6**). Here, we chose the final values that gave the lowest cost to use in the final parameter set.

### Model validation with experimental data measuring the impact of soluble BCMA on ADC efficacy

Elevated levels of soluble BCMA are frequently found in patients with multiple myeloma, and have been considered a potential hindrance to the performance of ADCs targeting membrane-bound BCMA [27,43]. Thus, we validated our model against a separate experimental dataset by running simulations to predict cell survival over various doses and at different levels of soluble BCMA. Simulations show that addition of soluble BCMA to the NCI-H929 cell culture system (at 0, 75, 270, and 720 ng/mL) does not cause changes to cell survival, which is consistent with the experimental data (**Figure 3D**). As the anti-BCMA ADC has stronger binding to membrane-bound BCMA than to soluble BCMA, this likely enables the ADC to overcome the effects of excess soluble BCMA in the system.

### Sensitivity Analysis

We performed both local and global sensitivity analyses on the parameters to explore the impact of these perturbations on key model outputs. First, we conducted a univariate sensitivity analysis to examine different outputs, looking at the effect on the area under the curve (AUC) for the concentrations of extracellular warhead, intracellular warhead, warhead-DNA complex, DNA crosslinking, and the cell population (**Figure 4A**). The DNA crosslinking parameters (n and K_A_) had the largest impact on both DNA crosslinking and cell population, while the cell killing rate (k_kill_) also had a significant impact on the cell population. Of the other parameters, various volumes within the *in vitro* system, DAR, and antigen production rate were found to have notable effects on the system outputs. The sensitivity to various parameters generally strengthened (becoming more positive or more negative) over time following ADC exposure (**Figure 4B**). For most parameters the sensitivity also intensified with increasing receptor expression levels (**Figure 4C**) and ADC doses (**Figure 4D**), though exhibiting peak sensitivity at a given expression level and decreasing again above that.

**Figure 4.**
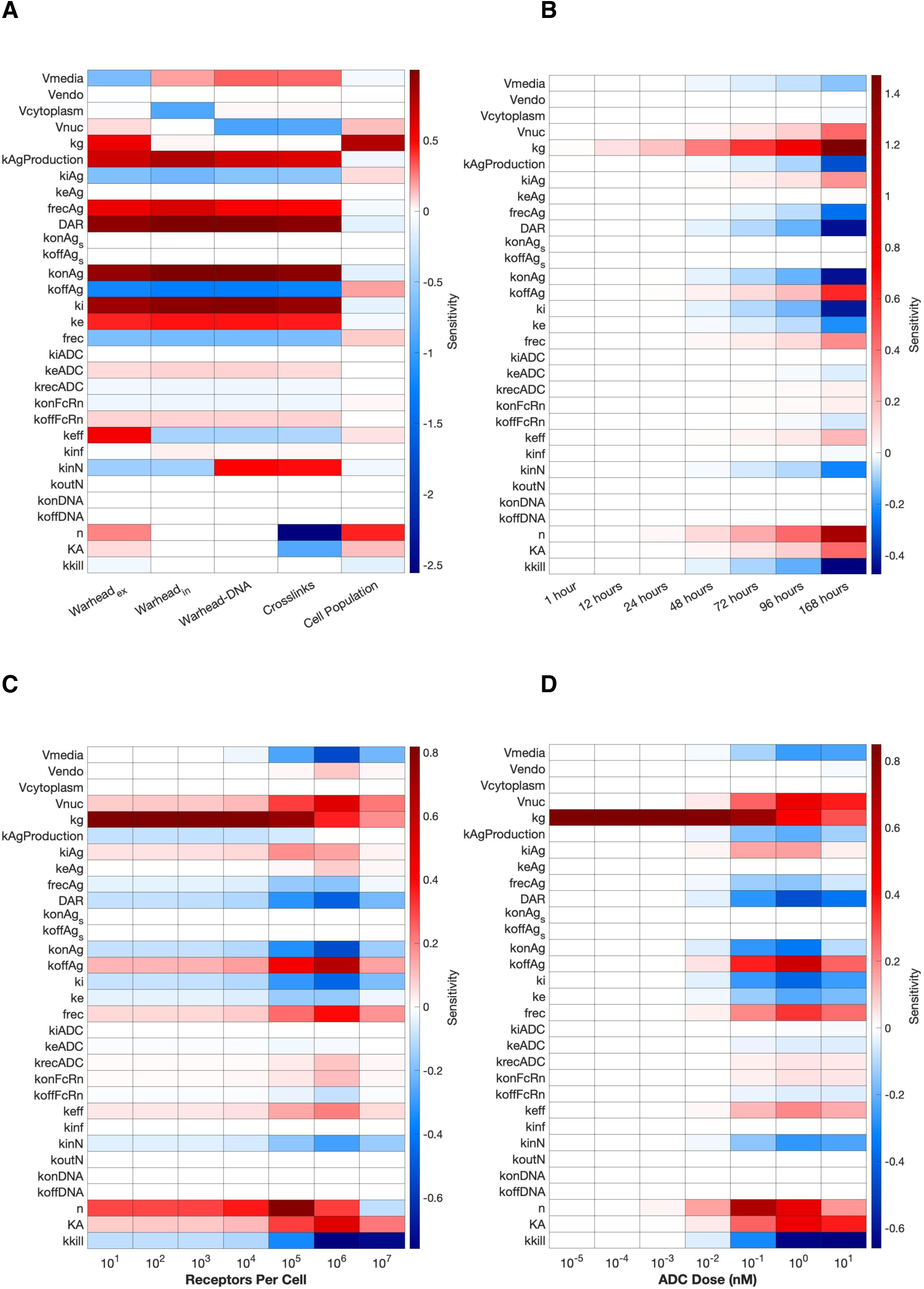
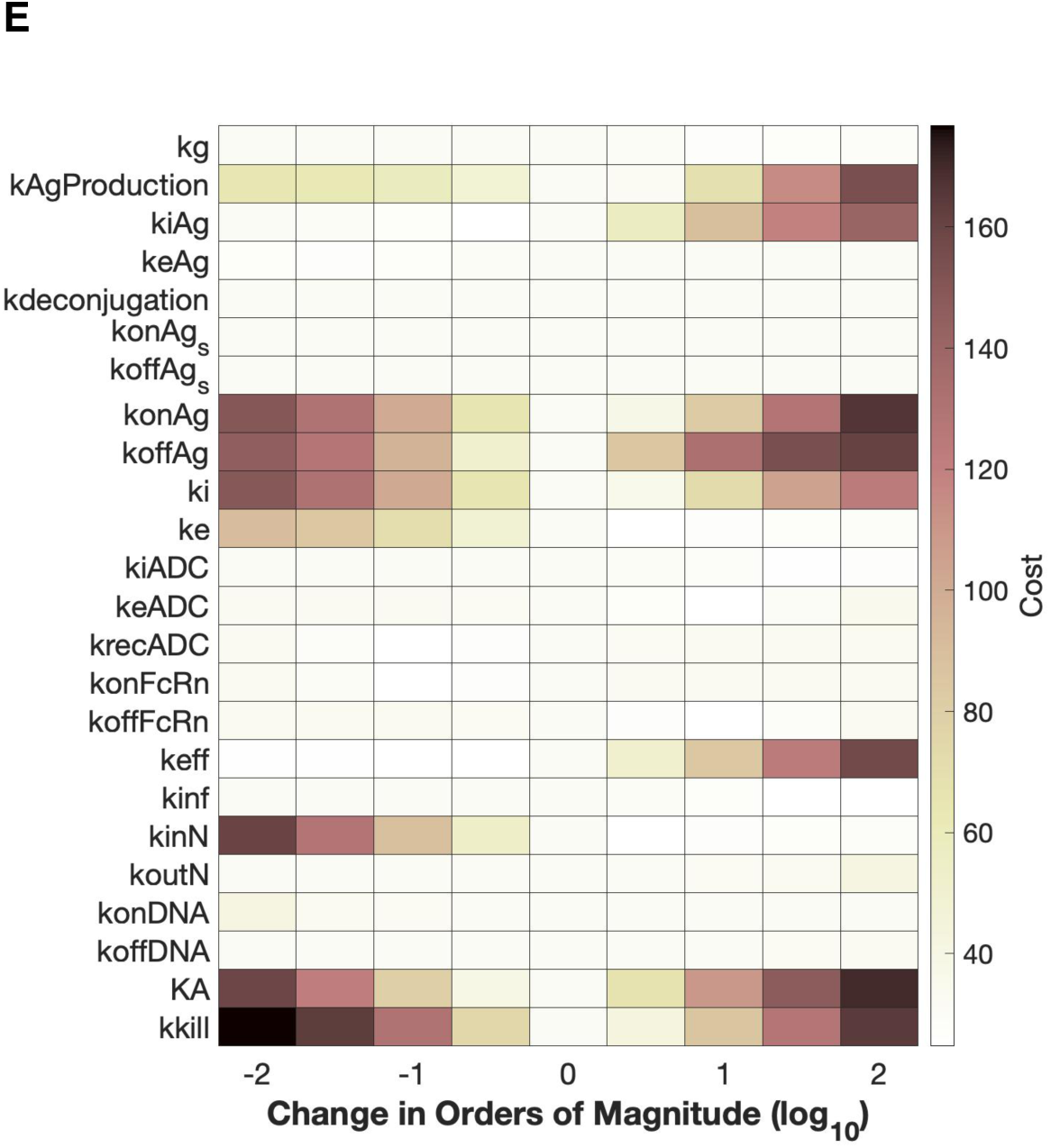
Sensitivity analyses for computational model parameters. **A**, Univariate local sensitivity analysis; we perturbed each parameter (y-axis) and examined the resulting sensitivity, defined as the percent change in the area under the curve (AUC) of each of the five key outputs (x-axis) per percent change in parameter. The outputs are (left to right) extracellular warhead, intracellular warhead, warhead bound to DNA, DNA crosslinking, and cell population. **B-D,** The local univariate sensitivity of cell population to the various parameters is different at different time points from 1 hour and 168 hours (7 days) following addition of the ADC **(B),** at different receptor expression levels from 101 to 107 receptors per cell **(C)**, and at different doses of MEDI2228 between 10-5 to 101 nM **(D)**. **E,** For a global sensitivity analysis of cell survival (ratio of treated to untreated cells) to the various parameters, parameter values were varied by up to two orders of magnitude above and below the original values, with the associated costs (root sum of squares error calculated against MEDI2228 cytotoxicity dose response experimental data) displayed on the heatmap. Unless otherwise noted, the simulations were run for 96 hours with an ADC dose of 4.1 x 10-2 nM and 1.9 x 104 receptors per cell.

We also performed a global sensitivity analysis to capture the effects over a wider range of parameter values (**Figure 4E**). The resulting heatmap shows the values of the costs calculated using the *rssq* function in MATLAB (the root sum of squares of the residuals between simulated and experimental *in vitro* cytotoxicity at 96 hours following MEDI2228 dosing) associated with modifying each parameter value over different orders of magnitude; the final selected parameter set aligns with those showing the lowest costs on the heatmap (center column). When varied over a few orders of magnitude, parameters involving antigen production, binding and unbinding of the ADC to Ag, warhead efflux, DNA crosslinking, and cell killing cause the largest shift in the calculated cost.

### Determining the ideal Drug-to-Antibody Ratio to maximize efficacy and minimize toxicity

Between the antibody, linker, and warhead components, ADCs have multiple design levers that can be manipulated to achieve differing outcomes. Here, we simulate modifications of these design levers to see how these changes can affect the ADC’s impact on cell killing. First, the number of warheads per antibody (Drug-to-Antibody Ratio, or DAR) is a key property in ADC design that can impact both efficacy and toxicity due to cell killing ability of the warhead. We simulated typical DAR values between 1 and 8 using the computational model to examine the effects on predicted *in vitro* cell survival and influxed warhead concentration in NCI-H929 cells incubated with ADC for 96 hours (**Figure 5**). The biggest jump in the rate and degree of cell killing occurs between DAR 1 and 2, while each additional warhead molecule added leads to smaller gains in the speed at which the cell population decreases and the ending cell survival (**Figure 5A**). The IC_50_s of the ADC with DAR 1 and DAR 8 differ by approximately 1 order of magnitude (**Figure 5C**). Depending on the dose of ADC given, the DAR value may not significantly impact the ending cell survival after 4 days (**Figure 5D**). Though this is not explored in this model, it is important to note that adding warheads to an ADC increases the size and bulk of the molecule, which may alter the ADC’s PK characteristics and lead to faster plasma clearance *in vivo* and lower stability [44,45]. Thus, additional DAR beyond 2 may not be worthwhile. It is also notable that the largest increase in concentration of influxed warhead (representative of bystander effects) also occurs between DAR 1 and 2 (**Figure 5B**). Depending on the situation, increasing the bystander effect may be a negative or a positive therapeutically, but if a negative then using DAR to increase efficacy is balanced by increased killing of nontarget cells.

**Figure 5.**
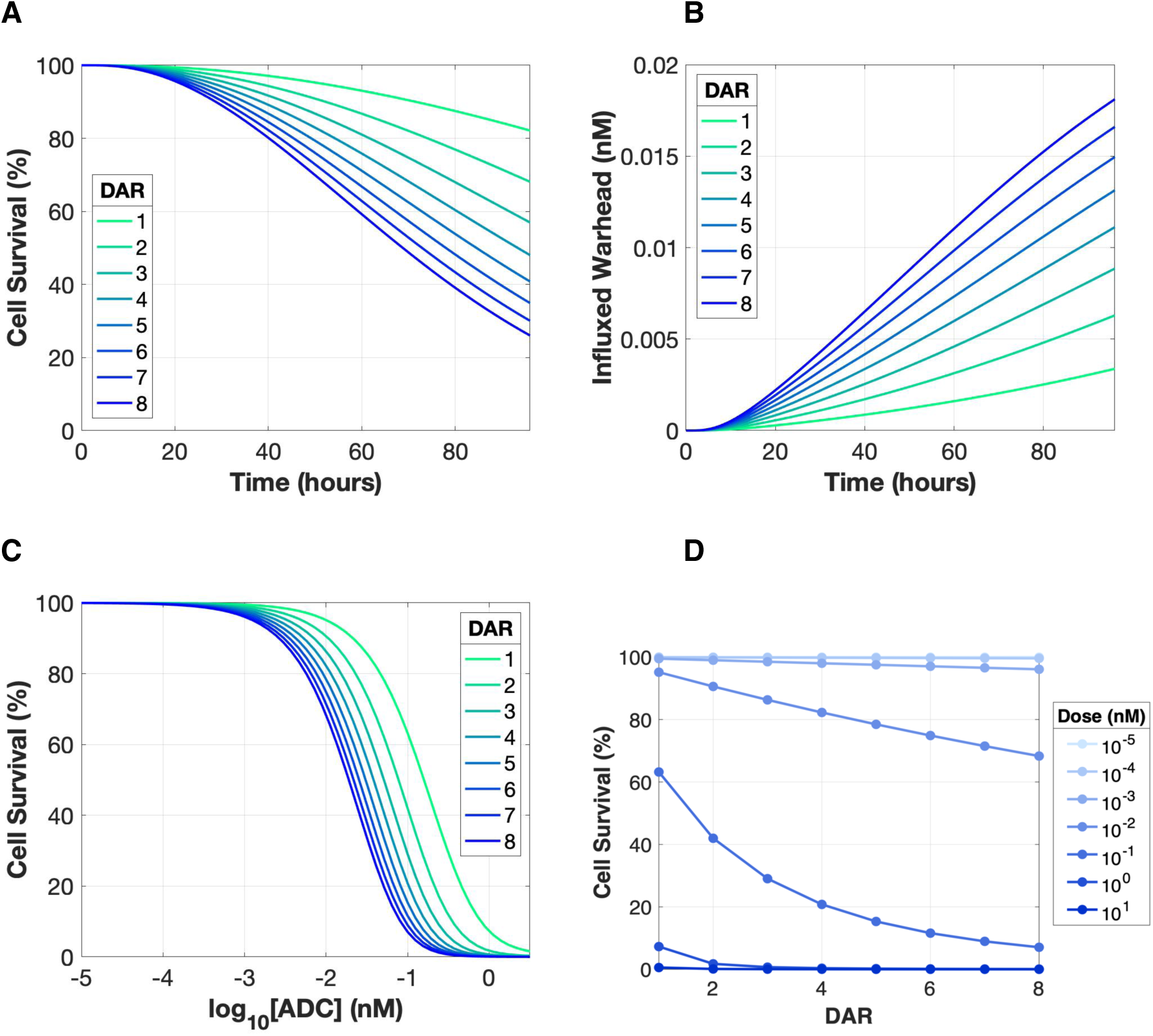
Increased Drug-to-Antibody Ratio (DAR) can increase warhead levels and cell killing, but may also increase bystander effects. **A-B**, we vary DAR over a range of 1 to 8 and simulate the predicted cell survival **(A)** and concentration of influxed warhead **(B)** over time following exposure to 4.1 x 10-2 nM ADC. While increasing DAR leads to gains in the degree and rate of cell killing, it also leads to greater levels of influxed warhead and thus potential bystander effects. **C-D**, Exploring dose dependency of ADC effect, we simulated cell survival versus dose at different DAR values **(C)** and cell survival versus DAR at different doses. **(D)** Depending on the dose of ADC used, increasing DAR also may not significantly increase cell killing.

### Warhead Potency and Lipophilicity

The therapeutic (and toxic) effects of the warhead are a combination of potency and availability. We simulated a wide range of parameter values for both the cell killing rate constant (k_kill_, 10^-3^ – 10^2^ hr^-1^) and the partition coefficient between intracellular and extracellular warhead (R, 10^-1^ – 10^3^), and predicted the impact on cell survival (**Figure 6**). Not surprisingly, increasing k_kill_ results in more and faster cell killing over time (**Figure 6A**), with the cell survival dose response curves shifting left with each increasing order of magnitude (**Figure 6B**). The partition coefficient (R) between intracellular and extracellular warhead is related to the lipophilicity of the PBD warhead, which has an estimated logD value of 4.12 (ref [41]). While increasing R, which is inversely proportional to the warhead efflux rate, also increases the speed at which the cultured cells die, these gains are nonlinear, particularly seen in the smaller changes in simulated cell survival versus time and dose between R = 100 and R = 1000 (**Figure 6C-D**). At lower R values, influx and efflux rates are more balanced, and warhead can more easily escape from inside the cell, resulting in less influxed warhead (**Figure 6E**). Interestingly, we notice that after the initial increase in warhead influx with increasing R (decreasing efflux rates, so warhead can stay inside the cell), this is followed by a significant decrease in influxed warhead as R increases from 100 to 1000 (**Figure 6E**). We hypothesize that this is due to the further decrease in efflux rates, where so little warhead can escape the cell such that there is less warhead available for influx.

**Figure 6.**
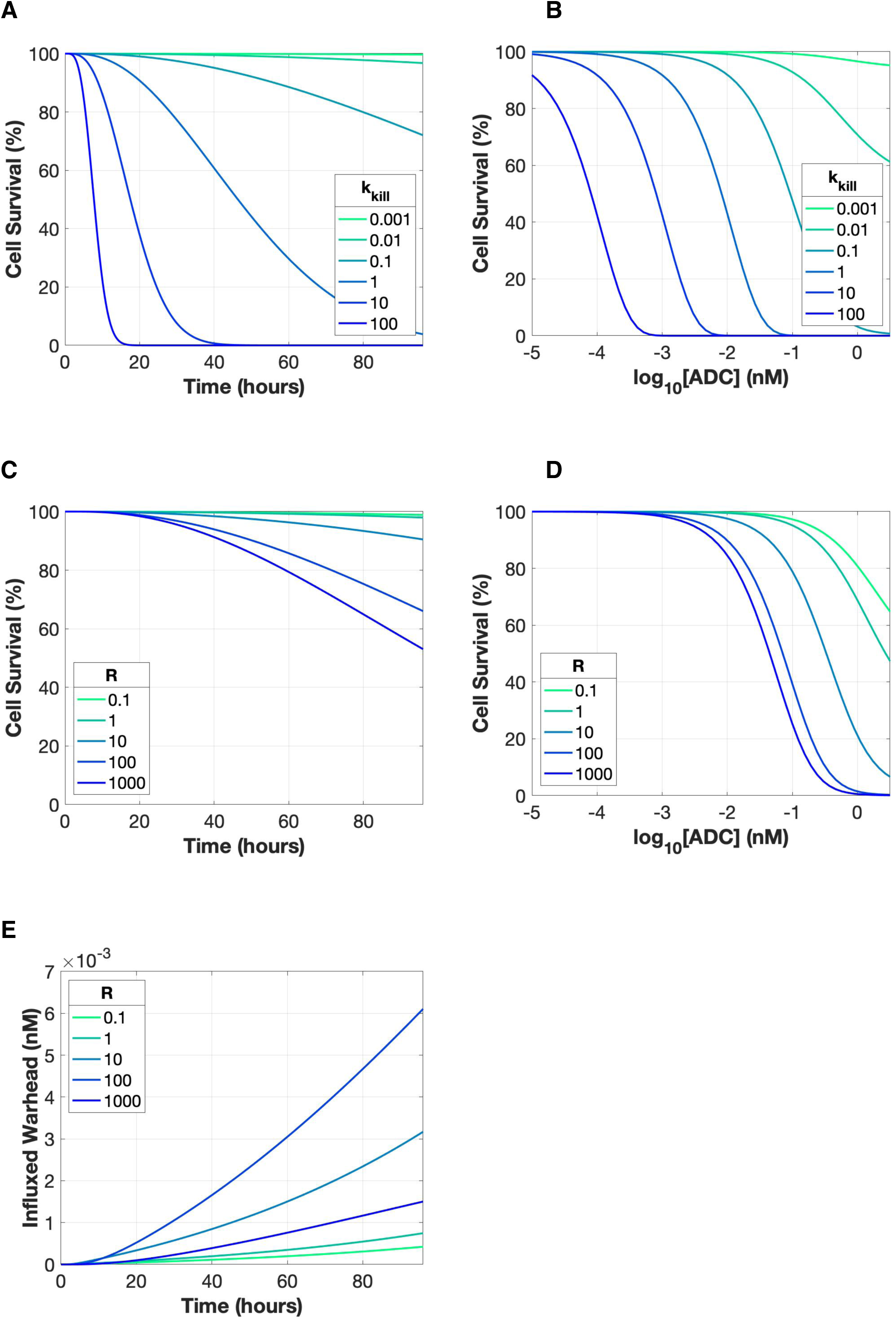
Warhead potency and lipophilicity can have stronger impact than DAR. **A-B**, We used the computational model to predict cell survival over time **(A)** and as a function of ADC dose **(B)**, at cell killing rate constants between 10-3 – 102 hr-1. **C-E**, We used the computational model to predict cell survival over time **(C)** and as a function of ADC dose **(D)**, and the concentration of influxed warhead over time **(E)** at values of the partition coefficient between intracellular and extracellular warhead (R) between 10-1 – 103. . Note that unlike DAR and kkill, response to R is not monotonic; after an initial increase in warhead with increasing R values, influxed warhead decreases substantially as R increases from 102 to 103.

### Tracking of warhead movement reveals the impact of linker design

The movement of warhead from inside the cell to outside and back is particularly relevant to the multicellular environment of tumors where the cell the warhead enters may not be the same type of cell that it left, i.e. not the type of cell targeted by the ADC, and the warhead can thus now kill the non-targeted cells; we explore that next.

Design of the linker is one of the cornerstones in ADC development, as the type of linker used can greatly impact how and where the warhead is released, and thus its ability to permeate into target or nearby cells. Understanding movement of the PBD warhead intracellularly and extracellularly can assist in elucidating the effects of linker design, which can be very difficult to explore experimentally. Therefore, we built tracking of different warhead populations into the computational model to computationally explore the impact of linker design. The model tracks the change in concentration over time for both extracellular and intracellular populations of warhead. Effectively, it tells us something about the recent history of the warhead. For extracellular warhead, there are two possible sources: warhead effluxed from inside the cell, and in the case of cleavable linkers, warhead deconjugated from ADC outside the cell. For intracellular warhead, there are four potential sources: warhead influx from outside the cell, which itself has two subtypes: warhead that was recently produced by extracellular ADC deconjugation (for cleavable linkers only), and warhead that was previously effluxed from cells; warhead that was recently produced from intracellular ADC cleavage in the same cell; and warhead from the nucleus (**Figure 7A**). Each of these populations are tracked separately within the model, enabling us to visualize how the warhead moves by seeing how these populations change over time.

**Figure 7.**
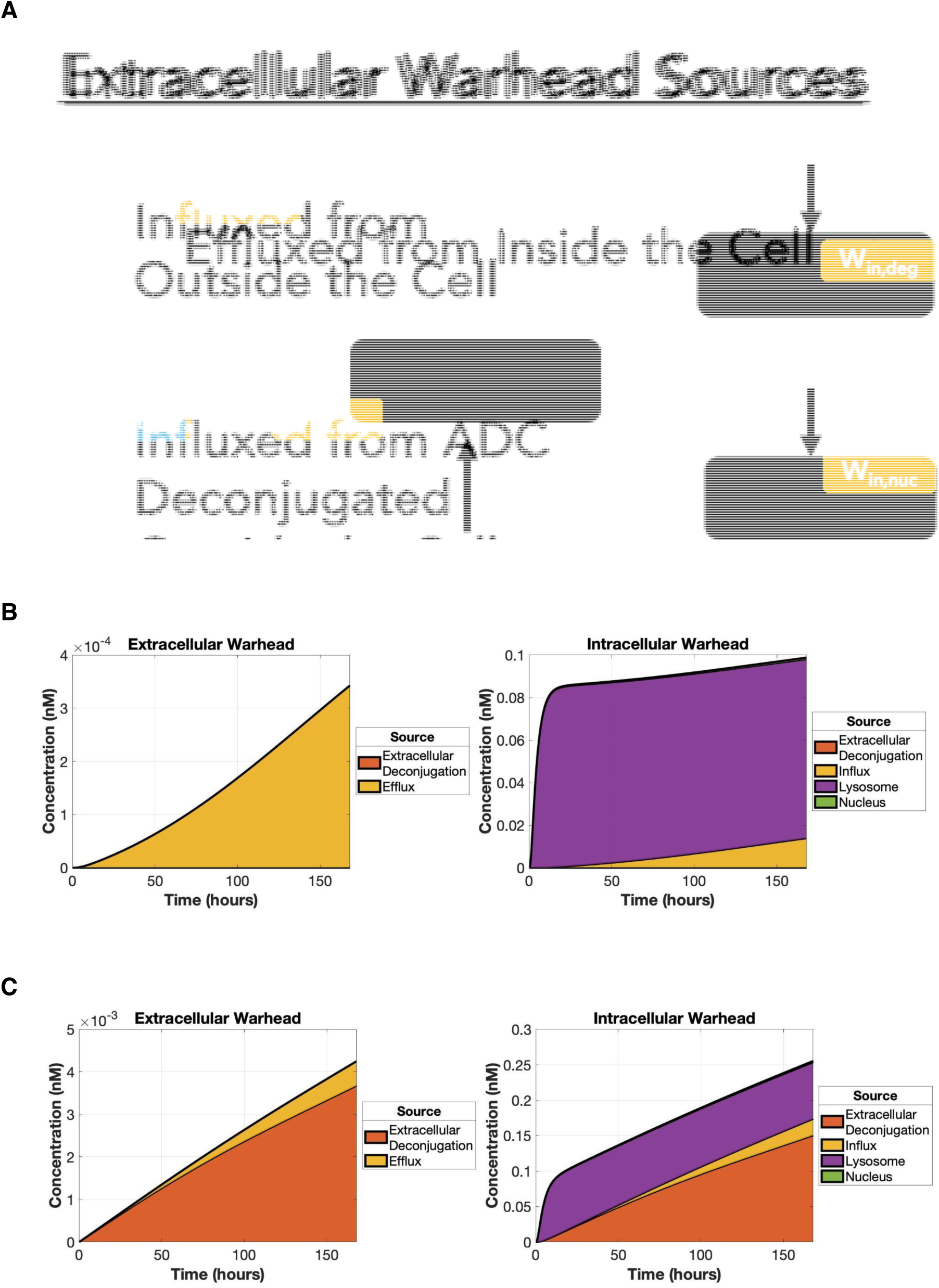
Tracking warhead location and recent history. Tracking populations of extracellular and intracellular warhead can reveal additional insights into warhead movement and mechanisms of action at the cellular and multicellular scales. **A,** There are several possible sources for both the extracellular and intracellular warhead subpopulations. **B,** For ADCs with noncleavable linkers, the external warhead population is entirely derived from efflux, while intracellular warhead comes from either lysosomal cleavage in that cell or influx of extracellular warhead previously cleaved in, and effluxed from, other cells. The impact of influx is limited under typical *in vitro* conditions where the volume of the media is much larger compared to the intracellular volume, due to dilution. **C,** In a hypothetical scenario for ADCs with cleavable linkers, extracellular warhead comes from both external deconjugation and efflux, while intracellular warhead comes from a combination of internal lysosomal cleavage, and two types of warhead coming from outside: influx of warhead that had previously come from another cell, and warhead that had most recently resulted from external deconjugation. There is also a small contribution of warhead returning from the nucleus. Depending on the rate of extracellular deconjugation, warhead influxed from extracellular deconjugation can become a strong contributor of the intracellular warhead levels.

Here, we simulate both noncleavable linkers (**Figure 7B**) and cleavable linkers (**Figure 7C**), which can be differentiated by the resulting warhead populations. Of note, MEDI2228 uses protease-cleavable linkers which are designed to degrade in the lysosome and remain stable in cell culture media; we chose to represent this by setting the rate of extracellular deconjugation to zero (similar to noncleavable linkers) for most simulations using this *in vitro* model. However, we also used serum stability data to optimize for the extracellular deconjugation rate constant and used this to explore the effects of warhead deconjugation outside the cell (to mimic cleavable linkers) in a few specific cases (**Figure S7**). In the case of noncleavable linkers, any extracellular warhead can only have come from inside the cell (W_ex,efflux_); for cleavable linkers, extracellular warhead can also originate from inside cells via efflux or can be deconjugated from ADC outside cells (W_ex,deg_). Similarly, for both types of linkers, a portion of warhead inside the cell was, at some point, outside the cell (W_in,influx_ and W_in,deg_), suggesting that bystander effects should be taken into account in both cleavable and noncleavable linkers for PBD ADCs. *In vivo*, the bystander effect might enhance therapeutic efficacy by increasing warhead concentration in target cells but can also contribute to the killing of nontarget cells; we explore this further in the following section.

### Effects of extracellular volumes in cell culture versus physiological conditions

Under cell culture conditions, the extracellular volume of the media is typically several orders of magnitude greater than volume inside the cells (e.g. typical 0.1 mL cell culture media vs 10^5^ cells * 1 pL/cell = 10^-4^ mL). However, under physiological conditions, the extracellular volume of the interstitial space is comparable to the volume inside the cells. Obviously, the total interstitial volumes *in vivo* can be large, but so too is the number of cells; for example, in a typical tumor, the intracellular volume may be 60% of the tissue volume while the interstitial space accounts for about 20-40% [46,47]. Another way of viewing this difference would be the surface-area-to-volume (SAV) ratio (i.e. surface area of cells vs extracellular volume; this is relevant to the interaction of antibodies with surface antigens); for a given number of cells, the *in vitro* cell culture has a much lower cell SAV ratio than *in vivo*. Thus, we ran simulations over a range of extracellular volumes (between 10^-8^ and 10^-4^ L) to simulate what happens at different SAV ratios, including levels typical of *in vitro* and *in vivo* conditions. Our simulations demonstrate that at lower extracellular volumes (similar to physiological conditions), a larger proportion of the total cytosolic warhead concentration comes from influxed warhead (**Figure 8**), suggesting that cells are more susceptible to bystander effects under physiological conditions than at typical cell culture conditions.

**Figure 8.**
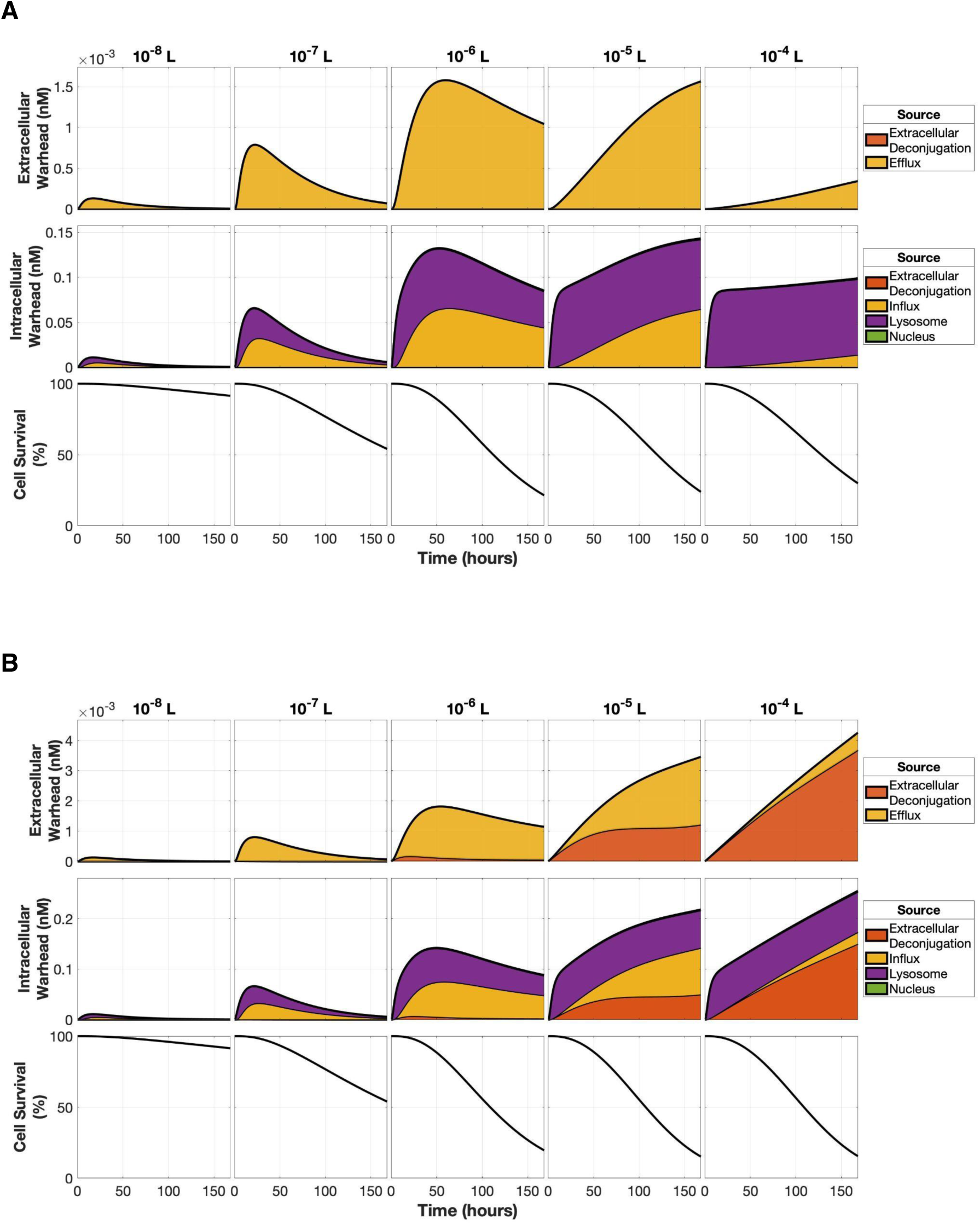
Extracellular media volume impacts ADC efficacy and bystander effects. Plots show extracellular and intracellular warhead concentrations (and tracked warhead subpopulations), plus cell survival at different extracellular media volumes (from 10-8 to 10-4), at a consistent initial extracellular ADC concentration of 4.1 x 10-2 nM. **A,** For ADCs with noncleavable linkers, extracellular warhead is entirely due to efflux; at lower volumes, there is less dilution but at very low volumes there is less total ADC which is consumed quickly. Intracellular warhead follows a similar overall pattern with volume, however as the extracellular levels get higher a larger proportion of the intracellular warhead comes from influx. Cell survival is dependent on overall intracellular warhead, rather than the specific source. **B**, For ADCs with cleavable linkers, similar trends can be seen, with an increasing proportion of extracellular warhead having been effluxed and an increasing proportion of intracellular warhead having been influxed (representative of the bystander effect) as volumes decrease. Extracellular deconjugation is a major contributor of warhead at higher extracellular volumes, but is limited at lower volumes as there is less total ADC in the system.

### Manipulating ADC design levers simultaneously to evaluate efficacy and potential toxicity

Now that we can see that bystander effects would likely be relevant *in vivo*, we return to considering the balance of ADC design considerations. We varied three mechanistic design parameters: DAR; warhead potency (k_kill_); and warhead lipophilicity (R, the partition coefficient between intracellular and extracellular warhead), to evaluate the effects on key outputs representing efficacy and toxicity (**Figure 9**). Though ADCs with the highest DAR and warhead potency enable the fastest killing of the cancer cell population (lower right corners of first row), they also demonstrate the most potential bystander effects, particularly when warhead potency is taken into account (lower right corners of third row). This effect is diminished at higher R values, which represent more lipophilic warheads that are sequestered into cells and show little or no bystander effects. Similar to what was seen in **Figure 6E**, we see a reversal in the amount of influxed warhead as R increases from 100 to 1000. Depending on whether or not bystander effects are desired (e.g. using them to facilitate deeper penetration of drug into solid tumors), manipulating the combination of these parameters can be used to design the optimal ADC for a particular system.

**Figure 9.**
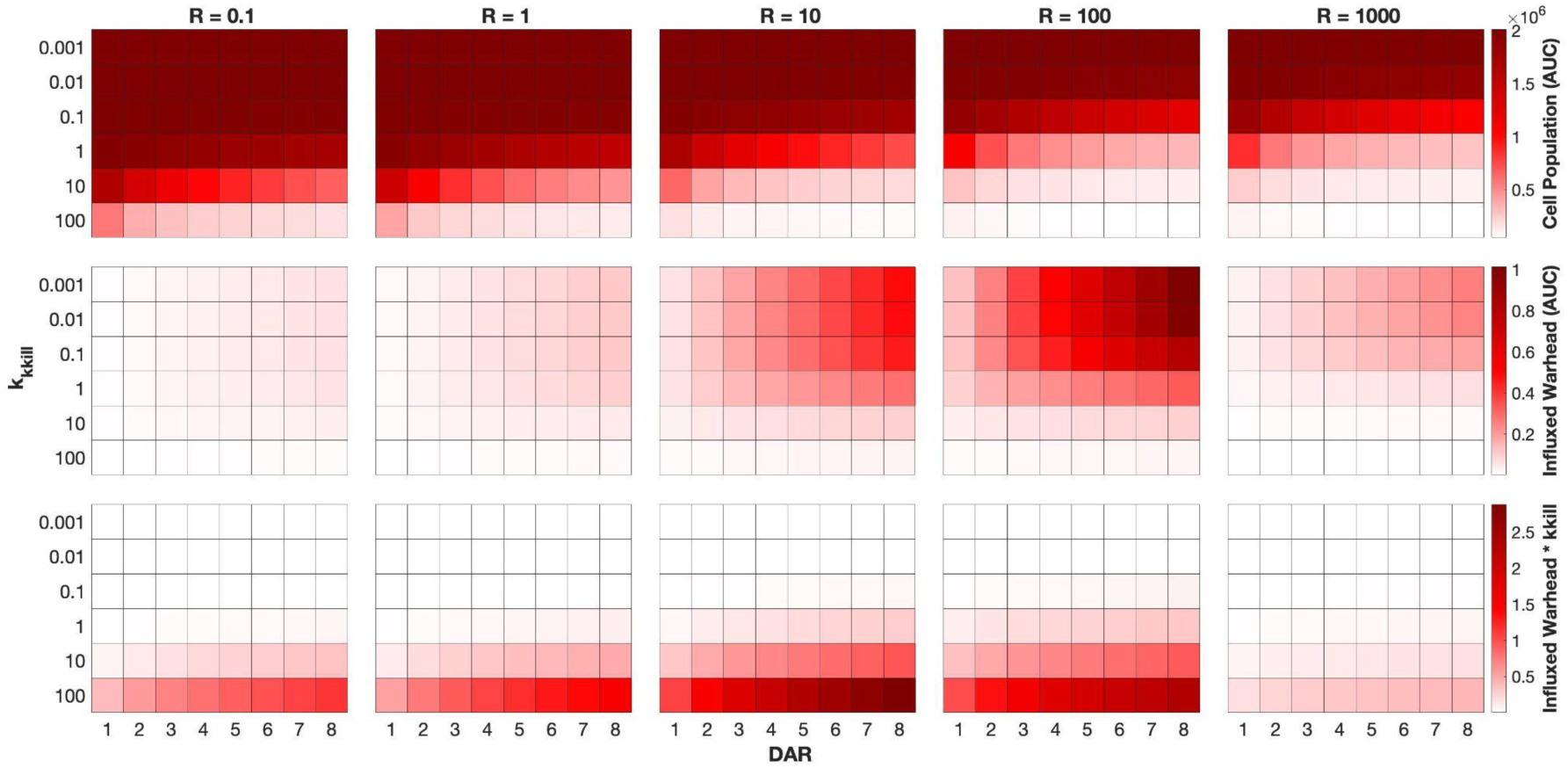
Impact of key ADC design parameters on efficacy and toxicity. To explore the role of key ADC design parameters on efficacy and toxicity *in vitro*, DAR values are varied between 1 to 8 (x-axis of each heatmap), kkill (warhead potency) values are varied between 10-3 to 102 (y-axis of each heatmap), and R (partition coefficient between intracellular and extracellular warhead) is varied between 10-1 to 103 (columns). Colors on each row indicate the resulting outputs: AUC of cell population (top) representing efficacy - how much and how quickly the cell population is killed by the ADC, with lighter colors indicating more and faster cell killing; AUC of influxed warhead (middle) could be used as an indicator of bystander effects, but is insufficient as it does not take into account the potency of the influxed warhead; and AUC of influxed warhead times kkill (bottom) which better represents the potential bystander killing ability of the influxed warhead. Using the model, these ADC design parameters (DAR, potency, and lipophilicity) can be altered to achieve the optimal combination of cell killing and bystander effects needed for the given conditions.

## Discussion

In this study, we have developed, parameterized, and validated a mechanistic model of antibody drug conjugates (ADCs), using in particular experimental data for MEDI2228, a BCMA-targeting antibody conjugated to the PBD warhead SG3199. While MEDI2228 demonstrated high preclinical efficacy in both cell culture and murine experiments along with relatively low toxicity within *in vivo* mouse models [27], concerns with efficacy and safety in humans were cited as the reasons for the termination of further clinical development after the phase 1 clinical trials [24,26,48,49]. Though MEDI2228 is no longer being actively developed, there is still much to learn from modeling therapies that do not succeed, particularly in understanding the reasons for the lack of success and translating this to design principles for future treatments. This is the first mechanistic model of a PBD-warhead ADC and of an anti-BCMA ADC, though the methodology can be generalized to other antibodies and other warheads.

We used this model to understand the impact of key ADC design considerations on both the direct efficacy and the potential toxicity (specifically, the magnitude of potential bystander effect) of the drug in *in vitro* cell culture. We designed the model equations to incorporate a tracking method so that we could estimate bystander effect size by quantifying the different recent sources of warhead in each location. By running the simulations with a range of surface-area-to-volume ratios, we also consider the likely implications of ADC design considerations on efficacy and toxicity *in vivo* and show in particular how the predicted behavior *in vitro* and *in vivo* can be different.

This computational model enables a detailed exploration of the biological processes governing ADC performance at the cellular scale. As the model can track the impact of both cell proliferation and cell killing, we can use simulations to compare the results with and without treatment, which is key to predicting cell survival (ratio of treated to untreated cells), which is the metric used in most of the experimental data. For instance, the sensitivity analysis shows that in addition to the expected importance of parameters such as antigen binding, warhead influx and efflux, and cell killing, other processes such as non-specific uptake and DNA crosslinking may also heavily influence cancer cell survival. Additionally, standard design levers of ADCs, including DAR, warhead potency, and warhead lipophilicity can be perturbed in the model in order to find the optimal combination of design features. Depending on the desired level of bystander killing, these parameters can be tailored to the specific cancer cell type and system being studied. Whether or not bystander killing is beneficial or toxic is likely a function of the cells being killed. In a microenvironment of mostly tumor cells, local efflux and then influx of warhead into another cell is most likely to increase warhead levels (and thus killing) of targeted tumor cells. If, however, there are cells that would be better spared - e.g. parenchymal tissue or stromal immune cells - then the bystander effect may cause the death of those cells.

Notably, we developed this computational model in such a way that it enables us to differentiate those on-target and off-target effects of the ADC and warhead. The warhead tracking feature built into the model allows us to predict both extracellular and intracellular concentrations of warhead over the course of a simulated experiment, which would typically very difficult to measure experimentally. It also allows us to predict the recent sources of those warhead subpopulations, which would likely be completely infeasible experimentally. This allows us to hone in on warhead movement within cells and between cells, and to better capture potential bystander effects, enabling us to compare the effect of warhead potency on both efficacy and potential toxicity.

In terms of ADC design considerations, we explored different values of warhead density (DAR), warhead potency (k_kill_) and warhead lipophilicity (R). The simulations show that increasing both the strength and availability of the warhead increases cell killing and therefore the efficacy of the ADC. However, there is a concomitant increase in the predicted bystander effect. Therefore, the killing of non-targeted cells likely needs to be considered. Of particular note, when we use the model to simulate *in vitro* cell culture, the magnitude of the bystander effect (i.e. the amount of warhead influxed from other cells) is low. However, we show here that this is largely a result of the large media reservoir outside the cells diluting the warhead. When we simulated lower extracellular volumes, more closely representing the surface-area-to-volume ratios seen *in vivo*, we can see (a) higher levels of intracellular warhead and (b) a much larger proportion of this being due to influx from other cells.

Construction of quantitative systems pharmacology models relies heavily on the experimental data available to parameterize the model. To maximize the existing data for anti-BCMA PBD ADCs to describe the underlying biological mechanisms, we used a sequential optimization process to isolate certain mechanisms and fit the relevant parameters to the data from various experiments. Biologically, the mechanism of ADC action typically starts from ADC binding to Ag, followed by internalization and then release of warhead. However, identifying model parameters with a “bottom-up” approach (beginning with the warhead-dependent mechanisms that ultimately determine the efficacy of the ADC at the single cell level) helps us to isolate and identify parameter values for that smaller core system describing warhead mechanisms of action, parameter values that should still hold when simulating within the framework of the whole ADC system. By contrast, optimization of the ADC/Ab-dependent parameters without consideration of the downstream warhead parameters could have led to parameter sets that do not fully represent the underlying biology during warhead-only simulations. The sequential optimization process also reduces the number of parameters being optimized at a given time, improving computational efficiency and the likelihood of consistent results.

As *in vitro* cytotoxicity assays are commonly performed when evaluating an ADC therapeutic candidate, we used data from different types of cytotoxicity experiments to parameterize the model, while supplementing with additional experimental data when possible. For instance, we optimized warhead-dependent parameters from warhead cytotoxicity data, and supplemented this with DNA-crosslinking dose response data to add greater detail on warhead mechanism of action. Thus, we are able to incorporate specific PBD-induced formation of DNA crosslinks, which have not previously been explored in QSP models of ADCs. This modular feature of the model can be applied to other ADC warheads that bind to DNA to induce DNA damage. For instance, calicheamicins (such as ozogamicin, which is currently used in 2 approved ADCs, gemtuzumab ozogamicin and inotuzumab ozogamicin) are a class of ADC warheads that also bind to the minor groove of DNA and cause DNA alkylation [50]; the DNA binding and fraction of alkylation can be described using this model by replacing fraction crosslinked for fraction alkylated.

Similarly, we found that isotype control ADC cytotoxicity data could be used to identify the non-specific uptake parameters in the model. While isotype control ADCs (which carry the same payload as the regular ADC but cannot bind to the target receptor) are typically used to compare and evaluate the effects of antibody binding experimentally, this data can also be incorporated into computational models, as the only method for these ADCs to enter the cell is via non-specific uptake. Further, we included experimentally-measured values for MEDI2228-FcRn binding and unbinding rate constants in the model. Ab-FcRn binding and recycling are important considerations in the development of antibody-based therapies such as ADCs, as FcRn binding and recycling contribute to the half-life of the ADC, and increased rates may prolong the half-life and reduce ADC accumulation in normal cells, thereby improving the therapeutic index of the ADC [40,51–53]. Mathematical models of FcRn binding have also been developed, and some ADC computational models have also incorporated FcRn mechanisms [54,42,55,37]. Here, we incorporate not only the FcRn recycling mechanism, but also ADC-FcRn binding and unbinding rate constants taken directly from experimental values, and modifications to FcRn can be easily altered in the model to determine the resulting effects. Creative use of existing data that are not typically used for modeling can enable us to identify parameters and add mechanistic detail into the model to more accurately depict the underlying biological processes.

We have incorporated modification terms to certain processes in the model to better represent the *in vitro* experimental system being simulated. For instance, while the influx of PBD involves movement through the phospholipid bilayer of the cell and is driven by the concentration and the hydrophobicity/hydrophilicity of the drug, ADC pinocytosis consists of an exchange of fluid between the extracellular and intracellular space and is volume-driven. Compared to *in vivo* systems where the volume differences between the extracellular and intracellular space are minimal, *in vitro* systems typically have a much higher extracellular to intracellular volume ratio due to the amount of media used for cell culture. Thus, these differences in volumes must be accounted for in order to prevent overrepresentation of molecules moving from the extracellular to intracellular compartments.

As described above, we have incorporated multiple biological mechanisms - some explored in previous models, some less so - into the present model. Although incorporating more biological mechanisms can allow for the creation of a more complete and descriptive model, efforts can be limited by the availability and type of supporting data. For instance, limited data was available on the kinetics of PBD-DNA binding, so we estimated the binding and unbinding rate constants for the SG3199 warhead based on the range of reaction rate constants for anthramycin, a precursor to PBD; these parameter values can be further refined when more experimental data becomes available. The model also includes a cell killing parameter which describes warhead potency rather than solely cell death, as it is not possible to distinguish cell killing from cell cycle arrest from these cytotoxicity assays. The model also does not account for DNA repair mechanisms that may reverse some of the damage caused by the formation of interstrand DNA crosslinks [29]. Additionally, active transport of PBD dimers via efflux pumps (particularly the membrane-bound drug transporter P-glycoprotein, which has been implicated in acquired small-molecule drug resistance for other ADCs) can be an important consideration when determining overall warhead efficacy and bystander effects [56,57], and may be incorporated into the model if the data are available.

We also assumed the model compartments were well-mixed, and that the rate of antigen production was constant, though this rate may be variable depending on the state of cells, as antigen production may be altered in response to ADC binding. Upon cell death, the warhead within the cell is not re-released into the system.

In summary, we have developed a computational mechanistic systems pharmacology model of the anti-BCMA PBD ADC MEDI2228 and applied it to simulate *in vitro* cell culture as well as modifying it to predict likely differences between *in vitro* and *in vivo* scenarios. This model includes both cellular and intracellular mechanisms specific to ADCs with PBD warheads and contains parameters from literature or fit to experimental data. The modularity of this model enables its use as a platform to describe other PBD ADCs. We have added a warhead tracking feature to the model which revealed the prevalence of bystander effects, which may be a source of unwanted toxicity. Using the relevant experimental data, this *in vitro* model can be expanded to incorporate whole-body pharmacokinetics, to serve as the basis for *in vivo* and clinical models of PBD ADCs. The resulting models will describe whole-body pharmacology and can be further developed to create a human clinical model, which can be used to run virtual clinical trials. The model and approach could also be generalized to other ADCs.

## Code Availability

All code written in support of this publication and relevant data are available at https://github.com/inezlam/pbd-adc-model-bcma-in-vitro. Code for the model simulations was written in MATLAB using version R2024a.

## References

1. Chabner BA, Roberts TG. Chemotherapy and the war on cancer. Nat Rev Cancer. 2005;5: 65–72. doi:10.1038/nrc1529

2. Galmarini D, Galmarini CM, Galmarini FC. Cancer chemotherapy: A critical analysis of its 60 years of history. Critical Reviews in Oncology/Hematology. 2012;84: 181–199. doi:10.1016/j.critrevonc.2012.03.002

3. Scialdone L. Overview of Supportive Care in Patients Receiving Chemotherapy: Antiemetics, Pain Management, Anemia, and Neutropenia. Journal of Pharmacy Practice. 2012;25: 209–221. doi:10.1177/0897190011431631

4. Chan A, Lees J, Keefe D. The changing paradigm for supportive care in cancer patients. Support Care Cancer. 2014;22: 1441–1445. doi:10.1007/s00520-014-2229-9

5. Remesh A. Toxicities of anticancer drugs and its management. Int J Basic Clin Pharmacol. 2012;1: 2. doi:10.5455/2319-2003.ijbcp000812

6. Dy GK, Adjei AA. Understanding, recognizing, and managing toxicities of targeted anticancer therapies. CA: A Cancer Journal for Clinicians. 2013;63: 249–279. doi:10.3322/caac.21184

7. Li J, Chen F, Cona MM, Feng Y, Himmelreich U, Oyen R, et al. A review on various targeted anticancer therapies. Targ Oncol. 2012;7: 69–85. doi:10.1007/s11523-012-0212-2

8. Schirrmacher V. From chemotherapy to biological therapy: A review of novel concepts to reduce the side effects of systemic cancer treatment (Review). International Journal of Oncology. 2019;54: 407–419. doi:10.3892/ijo.2018.4661

9. Scott AM, Allison JP, Wolchok JD. Monoclonal antibodies in cancer therapy. Cancer Immun. 2012;12: 14.

10. Zahavi D, Weiner L. Monoclonal Antibodies in Cancer Therapy. Antibodies. 2020;9: 34. doi:10.3390/antib9030034

11. Sievers EL, Senter PD. Antibody-drug conjugates in cancer therapy. Annual review of medicine. 2013;64: 15–29.

12. Peters C, Brown S. Antibody–drug conjugates as novel anti-cancer chemotherapeutics. Bioscience reports. 2015;35: e00225.

13. Thomas A, Teicher BA, Hassan R. Antibody–drug conjugates for cancer therapy. The Lancet Oncology. 2016;17: e254–e262.

14. Beck A, Goetsch L, Dumontet C, Corvaïa N. Strategies and challenges for the next generation of antibody–drug conjugates. Nat Rev Drug Discov. 2017;16: 315–337. doi:10.1038/nrd.2016.268

15. Fu Z, Li S, Han S, Shi C, Zhang Y. Antibody drug conjugate: the “biological missile” for targeted cancer therapy. Sig Transduct Target Ther. 2022;7: 1–25. doi:10.1038/s41392-022-00947-7

16. McCombs JR, Owen SC. Antibody Drug Conjugates: Design and Selection of Linker, Payload and Conjugation Chemistry. AAPS J. 2015;17: 339–351. doi:10.1208/s12248-014-9710-8

17. Gauzy-Lazo L, Sassoon I, Brun M-P. Advances in Antibody–Drug Conjugate Design: Current Clinical Landscape and Future Innovations. SLAS DISCOVERY: Advancing the Science of Drug Discovery. 2020;25: 843–868. doi:10.1177/2472555220912955

18. Samantasinghar A, Sunildutt NP, Ahmed F, Soomro AM, Salih ARC, Parihar P, et al. A comprehensive review of key factors affecting the efficacy of antibody drug conjugate. Biomedicine & Pharmacotherapy. 2023;161: 114408. doi:10.1016/j.biopha.2023.114408

19. Clegg LE, Mac Gabhann F. Molecular mechanism matters: Benefits of mechanistic computational models for drug development. Pharmacological research. 2015;99: 149–154.

20. Xie L, Draizen EJ, Bourne PE. Harnessing big data for systems pharmacology. Annual review of pharmacology and toxicology. 2017;57: 245–262.

21. van der Graaf PH, Benson N. Systems Pharmacology: Bridging Systems Biology and Pharmacokinetics-Pharmacodynamics (PKPD) in Drug Discovery and Development. Pharm Res. 2011;28: 1460–1464. doi:10.1007/s11095-011-0467-9

22. Knight-Schrijver VR, Chelliah V, Cucurull-Sanchez L, Le Novère N. The promises of quantitative systems pharmacology modelling for drug development. Computational and Structural Biotechnology Journal. 2016;14: 363–370. doi:10.1016/j.csbj.2016.09.002

23. Liu K, Li M, Li Y, Li Y, Chen Z, Tang Y, et al. A review of the clinical efficacy of FDA-approved antibody‒drug conjugates in human cancers. Molecular Cancer. 2024;23: 62. doi:10.1186/s12943-024-01963-7

24. Dumontet C, Reichert JM, Senter PD, Lambert JM, Beck A. Antibody–drug conjugates come of age in oncology. Nat Rev Drug Discov. 2023;22: 641–661. doi:10.1038/s41573-023-00709-2

25. Coats S, Williams M, Kebble B, Dixit R, Tseng L, Yao N-S, et al. Antibody–drug conjugates: future directions in clinical and translational strategies to improve the therapeutic index. Clinical Cancer Research. 2019;25: 5441–5448.

26. Maecker H, Jonnalagadda V, Bhakta S, Jammalamadaka V, Junutula JR. Exploration of the antibody–drug conjugate clinical landscape. MAbs. 2023;15: 2229101. doi:10.1080/19420862.2023.2229101

27. Kinneer K, Flynn M, Thomas SB, Meekin J, Varkey R, Xiao X, et al. Preclinical assessment of an antibody–PBD conjugate that targets BCMA on multiple myeloma and myeloma progenitor cells. Leukemia. 2019;33: 766–771. doi:10.1038/s41375-018-0278-7

28. Hartley JA. Antibody-drug conjugates (ADCs) delivering pyrrolobenzodiazepine (PBD) dimers for cancer therapy. Expert Opinion on Biological Therapy. 2021;21: 931–943. doi:10.1080/14712598.2020.1776255

29. Hartley JA, Flynn MJ, Bingham JP, Corbett S, Reinert H, Tiberghien A, et al. Pre-clinical pharmacology and mechanism of action of SG3199, the pyrrolobenzodiazepine (PBD) dimer warhead component of antibody-drug conjugate (ADC) payload tesirine. Scientific reports. 2018;8: 10479.

30. Kinneer K, Meekin J, Varkey R, Xiao X, Zhong H, Breen S, et al. Preclinical Evaluation of MEDI2228, a BCMA-Targeting Pyrrolobenzodiazepine-Linked Antibody Drug Conjugate for the Treatment of Multiple Myeloma. Blood. 2017;130: 3153. doi:10.1182/blood.V130.Suppl_1.3153.3153

31. Shah N, Chari A, Scott E, Mezzi K, Usmani SZ. B-cell maturation antigen (BCMA) in multiple myeloma: rationale for targeting and current therapeutic approaches. Leukemia. 2020;34: 985–1005. doi:10.1038/s41375-020-0734-z

32. Lam I, Pilla Reddy V, Ball K, Arends RH, Mac Gabhann F. Development of and insights from systems pharmacology models of antibody-drug conjugates. CPT: Pharmacometrics & Systems Pharmacology. 2022;11: 967–990.

33. Shah DK, Haddish-Berhane N, Betts A. Bench to bedside translation of antibody drug conjugates using a multiscale mechanistic PK/PD model: a case study with brentuximab-vedotin. Journal of pharmacokinetics and pharmacodynamics. 2012;39: 643–659.

34. Shah DK, King LE, Han X, Wentland J-A, Zhang Y, Lucas J, et al. A priori prediction of tumor payload concentrations: preclinical case study with an auristatin-based anti-5T4 antibody-drug conjugate. The AAPS journal. 2014;16: 452–463.

35. Maass KF, Kulkarni C, Betts AM, Wittrup KD. Determination of cellular processing rates for a trastuzumab-maytansinoid antibody-drug conjugate (ADC) highlights key parameters for ADC design. The AAPS journal. 2016;18: 635–646.

36. Vasalou C, Helmlinger G, Gomes B. A mechanistic tumor penetration model to guide antibody drug conjugate design. PloS one. 2015;10: e0118977.

37. Cilliers C, Guo H, Liao J, Christodolu N, Thurber GM. Multiscale modeling of antibody-drug conjugates: connecting tissue and cellular distribution to whole animal pharmacokinetics and potential implications for efficacy. The AAPS journal. 2016;18: 1117–1130.

38. Singh AP, Guo L, Verma A, Wong GG-L, Shah DK. A cell-level systems PK-PD model to characterize in vivo efficacy of ADCs. Pharmaceutics. 2019;11: 98.

39. Menezes B, Cilliers C, Wessler T, Thurber GM, Linderman JJ. An agent-based systems pharmacology model of the antibody-drug conjugate Kadcyla to predict efficacy of different dosing regimens. The AAPS journal. 2020;22: 1–13.

40. Mahalingaiah PK, Ciurlionis R, Durbin KR, Yeager RL, Philip BK, Bawa B, et al. Potential mechanisms of target-independent uptake and toxicity of antibody-drug conjugates. Pharmacology & Therapeutics. 2019;200: 110–125. doi:10.1016/j.pharmthera.2019.04.008

41. Khera E, Cilliers C, Bhatnagar S, Thurber GM. Computational transport analysis of antibody-drug conjugate bystander effects and payload tumoral distribution: implications for therapy. Molecular Systems Design & Engineering. 2018;3: 73–88.

42. Patsatzis DG, Wu S, Shah DK, Goussis DA. Algorithmic multiscale analysis for the FcRn mediated regulation of antibody PK in human. Sci Rep. 2022;12: 6208. doi:10.1038/s41598-022-09846-x

43. Sanchez E, Li M, Kitto A, Li J, Wang CS, Kirk DT, et al. Serum B-cell maturation antigen is elevated in multiple myeloma and correlates with disease status and survival. British Journal of Haematology. 2012;158: 727–738. doi:10.1111/j.1365-2141.2012.09241.x

44. Lyon R, Bovee T, Doronina S, Burke P, Hunter J, Neff-LaFord H, et al. Reducing hydrophobicity of homogeneous antibody-drug conjugates improves pharmacokinetics and therapeutic index. Nature biotechnology. 2015;33. doi:10.1038/nbt.3212

45. Sukumaran S, Zhang C, Leipold DD, Saad OM, Xu K, Gadkar K, et al. Development and translational application of an integrated, mechanistic model of antibody-drug conjugate pharmacokinetics. The AAPS journal. 2017;19: 130–140.

46. Madelin G, Kline R, Walvick R, Regatte RR. A method for estimating intracellular sodium concentration and extracellular volume fraction in brain in vivo using sodium magnetic resonance imaging. Sci Rep. 2014;4: 4763. doi:10.1038/srep04763

47. Lee JY, Ryu HS, Yoon SS, Kim EH, Yoon SW. Extracellular-to-Intracellular Fluid Volume Ratio as a Prognostic Factor for Survival in Patients With Metastatic Cancer. Integr Cancer Ther. 2019;18: 1534735419847285. doi:10.1177/1534735419847285

48. Ray U, Orlowski RZ. Antibody–Drug Conjugates for Multiple Myeloma: Just the Beginning, or the Beginning of the End? Pharmaceuticals (Basel). 2023;16: 590. doi:10.3390/ph16040590

49. MedImmune LLC. A Phase 1, Open-label Study to Evaluate the Safety, Pharmacokinetics, Immunogenicity, and Preliminary Efficacy of MEDI2228 in Subjects With Relapsed/Refractory Multiple Myeloma. clinicaltrials.gov; 2022 Mar. Report No.: NCT03489525. Available: https://clinicaltrials.gov/study/NCT03489525

50. Yaghoubi S, Karimi MH, Lotfinia M, Gharibi T, Mahi-Birjand M, Kavi E, et al. Potential drugs used in the antibody–drug conjugate (ADC) architecture for cancer therapy. J Cell Physiol. 2020;235: 31–64. doi:10.1002/jcp.28967

51. Ritchie M, Tchistiakova L, Scott N. Implications of receptor-mediated endocytosis and intracellular trafficking dynamics in the development of antibody drug conjugates. mAbs. 2013;5: 13–21. doi:10.4161/mabs.22854

52. Xu S. Internalization, Trafficking, Intracellular Processing and Actions of Antibody-Drug Conjugates. Pharm Res. 2015;32: 3577–3583. doi:10.1007/s11095-015-1729-8

53. Hamblett KJ, Le T, Rock BM, Rock DA, Siu S, Huard JN, et al. Altering Antibody–Drug Conjugate Binding to the Neonatal Fc Receptor Impacts Efficacy and Tolerability. Mol Pharmaceutics. 2016;13: 2387–2396. doi:10.1021/acs.molpharmaceut.6b00153

54. Ferl GZ, Wu AM, DiStefano JJ. A Predictive Model of Therapeutic Monoclonal Antibody Dynamics and Regulation by the Neonatal Fc Receptor (FcRn). Ann Biomed Eng. 2005;33: 1640–1652. doi:10.1007/s10439-005-7410-3

55. Shah DK, Betts AM. Towards a platform PBPK model to characterize the plasma and tissue disposition of monoclonal antibodies in preclinical species and human. J Pharmacokinet Pharmacodyn. 2012;39: 67–86. doi:10.1007/s10928-011-9232-2

56. Corbett S, Huang S, Zammarchi F, Howard PW, van Berkel PH, Hartley JA. The Role of Specific ATP-Binding Cassette Transporters in the Acquired Resistance to Pyrrolobenzodiazepine Dimer–Containing Antibody–Drug Conjugates. Molecular Cancer Therapeutics. 2020;19: 1856–1865. doi:10.1158/1535-7163.MCT-20-0222

57. Zammarchi F, Havenith KEG, Chivers S, Hogg P, Bertelli F, Tyrer P, et al. Preclinical Development of ADCT-601, a Novel Pyrrolobenzodiazepine Dimer-based Antibody–drug Conjugate Targeting AXL-expressing Cancers. Molecular Cancer Therapeutics. 2022;21: 582–593. doi:10.1158/1535-7163.MCT-21-0715

58. Sims CE, Allbritton NL. Analysis of single mammalian cells on-chip. Lab Chip. 2007;7: 423–440. doi:10.1039/B615235J

59. Heyden S, Ortiz M. Investigation of the influence of viscoelasticity on oncotripsy. Computer Methods in Applied Mechanics and Engineering. 2017;314: 314–322. doi:10.1016/j.cma.2016.08.026

60. Griffiths G, Back R, Marsh M. A quantitative analysis of the endocytic pathway in baby hamster kidney cells. J Cell Biol. 1989;109: 2703–2720. doi:10.1083/jcb.109.6.2703

61. Luzio JP, Pryor PR, Bright NA. Lysosomes: fusion and function. Nat Rev Mol Cell Biol. 2007;8: 622–632. doi:10.1038/nrm2217

62. Recombinant Human BCMA. In: PeproTech [Internet]. [cited 15 Feb 2024]. Available: https://www.peprotech.com/recombinant-human-bcma

63. Gazdar AF, Oie HK, Kirsch IR, Hollis GF. Establishment and characterization of a human plasma cell myeloma culture having a rearranged cellular myc proto-oncogene. Blood. 1986;67: 1542–1549.

