## Supplemental Information for "Quantitative systems pharmacology modeling of pyrrolobenzodiazepine antibody-drug conjugates targeting BCMA"

### Supplemental Figures

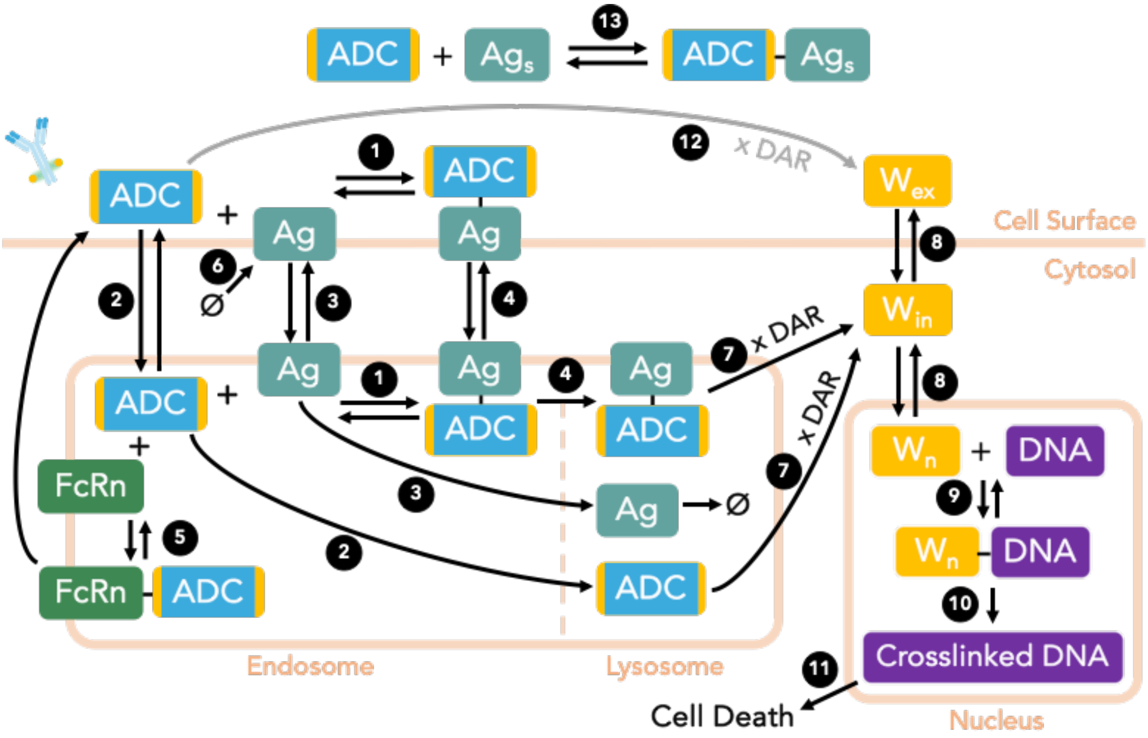


**Figure S1. Detailed Model Schematic.**
The model includes mechanisms of ADC binding and trafficking, warhead deconjugation and movement, and warhead mechanism of action. The antibody component of the ADC can bind reversibly to the antigen (1). ADC can also enter and leave the cell endosomes via pinocytosis (2), and this internal ADC can bind to endosomal antigen (1). The ADC-Ag surface complex can be internalized (4); the internal complex can be recycled back to the surface or sent to the lysosome irreversibly (4), as can the antigen and free ADC (3, 2). Endosomal ADC can also be recycled to the surface via binding to FcRn (5). In the lysosome, degradation of the ADC enables irreversible deconjugation of the warhead (7), which leaves to the cytoplasm. From here, the warhead can be effluxed from the cell (8) or enter the nucleus (8), both reversible processes. The nuclear warhead can bind reversibly to binding sites on DNA (9), and this complexes induces cross-linking of DNA (10) which induces cell death (11). Extracellular ADC may also deconjugate warhead if a suitable cleavable linker is used (12), and can bind to soluble, non-cell-associated antigen if present (13). Extracellular warhead, whether originating from extracellular deconjugation (12) or from efflux from a cell (8), can influx into cells as well (8).

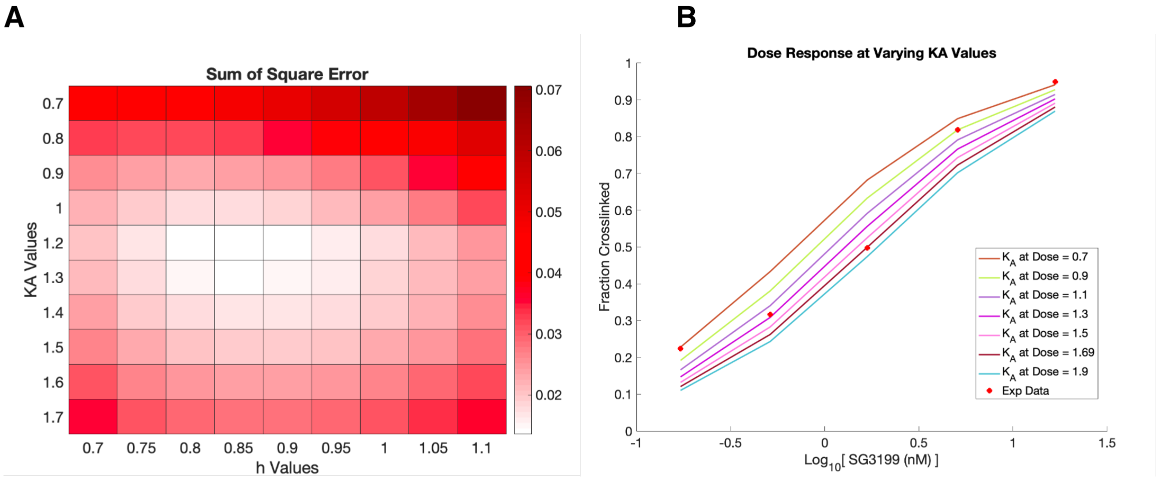


**Figure S2. Representing DNA Crosslinking using the Hill Equation.**
**A**, Heatmap of the total error (sum of squared differences between simulation and experimental results) over a range of possible values for the Hill parameters. **B**, Experimental data (dots) and simulation results (lines) for DNA crosslinking in response to different warhead doses, for different values of KA values while holding the Hill coefficient n constant.

**Figure S3. Optimization of 3 nonspecific uptake parameters to isotype control ADC cytotoxicity data.**

**A**, Starting with 100 different guesses across a wide range of values (x-axis) for each of the three parameters to be optimized - the internalization rate constant (pinocytosis), endosomal exit rate constant, and recycling rate constant of ADCs - we obtained optimized values that for lowest-cost fits were mostly not dependent on initial guess but only somewhat consistent/constrained. **B**, The simulation results (lines) for the lowest-cost optimization demonstrates good fit to the experimental data (dots).

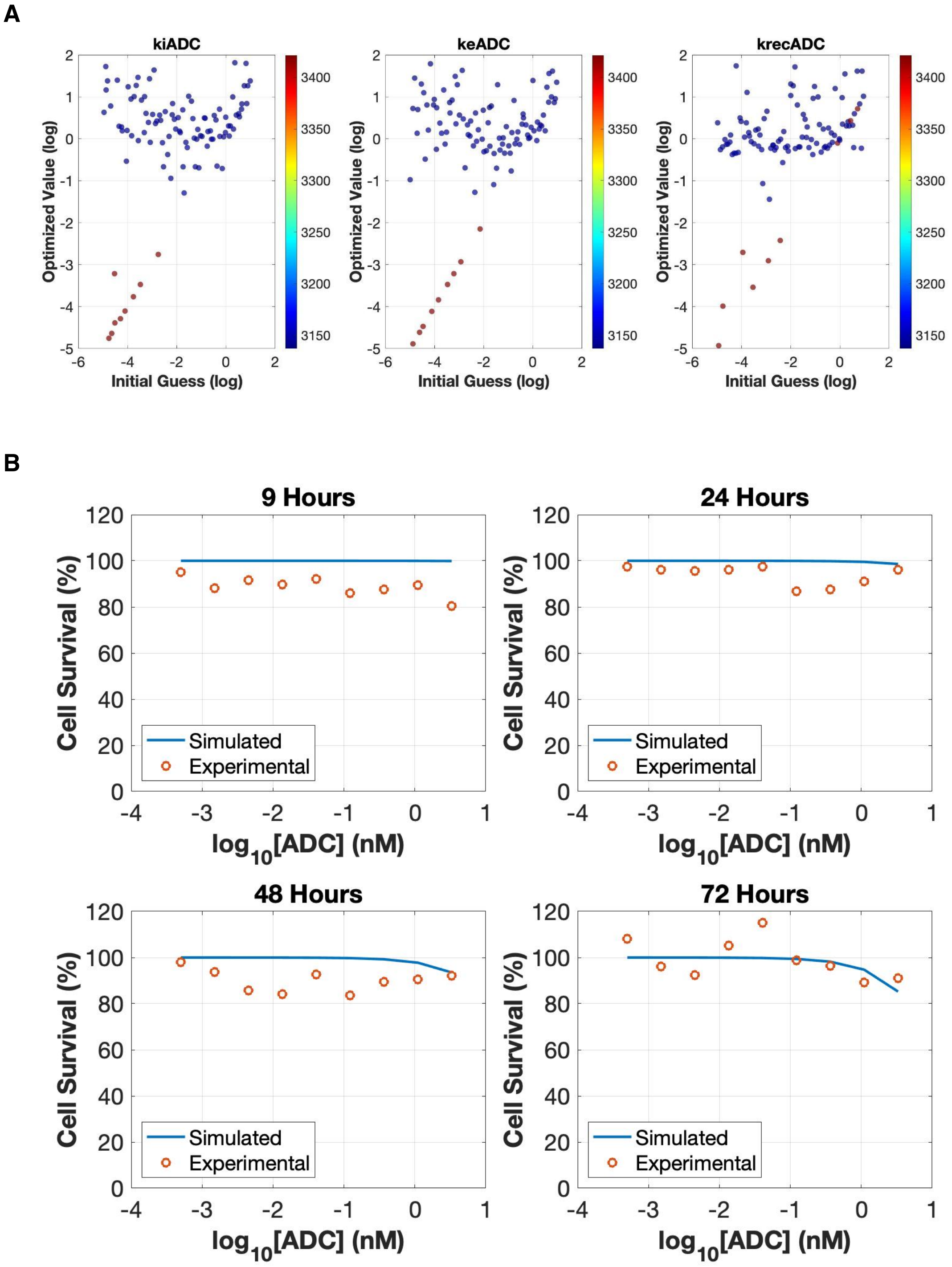
A
kiADC
keADC
krecADC
Optimized Value (log)
Initial Guess (log)
B
9 Hours
24 Hours
48 Hours
72 Hours
Cell Survival (%)
log10[ADC] (nM)
Simulated
Experimental

**Figure S4**. **Optimization of 2 nonspecific uptake parameters to isotype control ADC cytotoxicity data.**

**A**, Starting with 100 different guesses across a wide range of values (x-axis) for each of the two parameters to be optimized - the internalization rate constant (pinocytosis) and recycling rate constant of ADCs - we obtained optimized values that for lowest-cost fits were mostly not dependent on initial guess and while not unique were largely consistent and constrained. **B**, The simulation results (lines) for the lowest-cost optimization demonstrates good fit to the experimental data (dots).

A

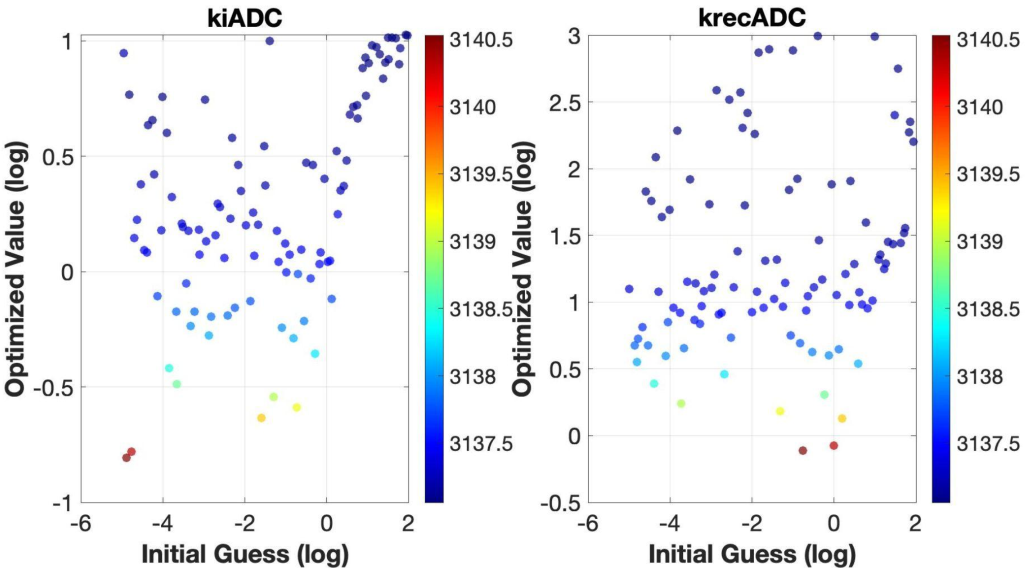
kiADC
krecADC
Optimized Value (log)
Initial Guess (log)
3140.5
3140
3139.5
3139
3138.5
3138
3137.5
1
0.5
0
-0.5
-1
-6
-4
-2
0
2
3
2.5
2
1.5
1
0.5
0
-0.5

B

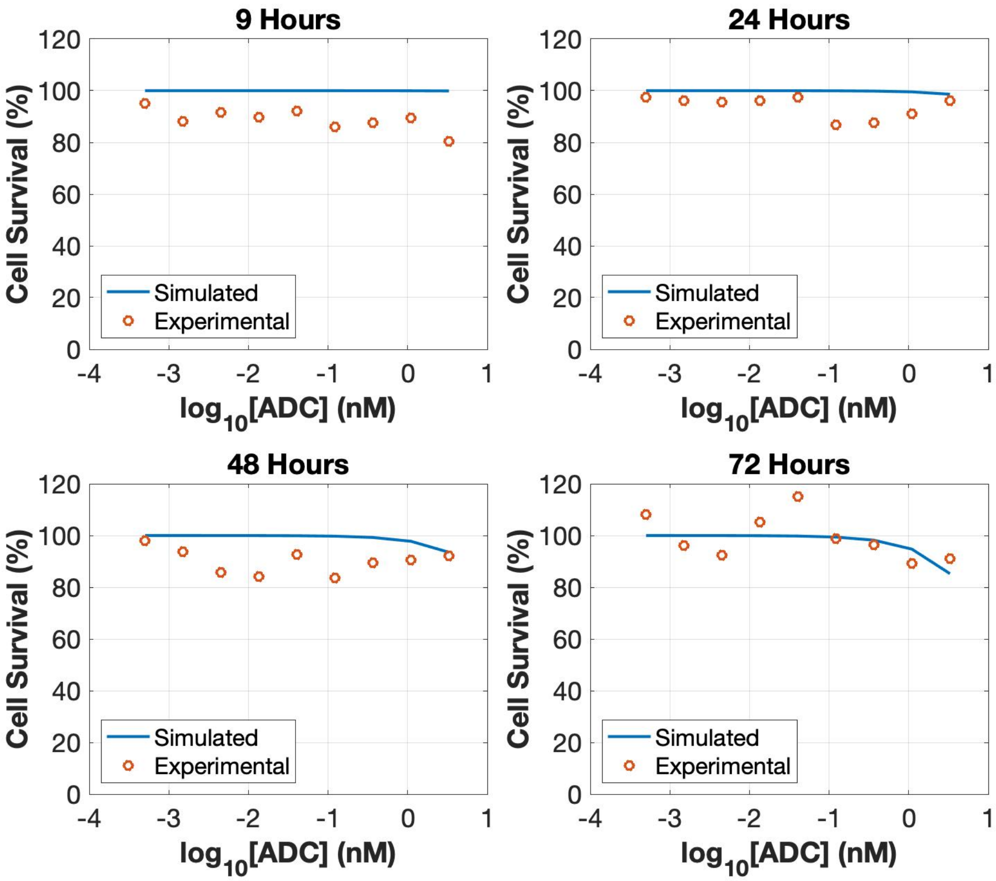
9 Hours
24 Hours
48 Hours
72 Hours
Cell Survival (%)
log10[ADC] (nM)
Simulated
Experimental
120
100
80
60
40
20
0
-4
-3
-2
-1
0
1

**Figure S5. Optimization of 6 receptor-mediated uptake parameters to ADC cytotoxicity data.** This includes nested optimization of antigen production rate to steady state levels in the absence of ADC. **A**, Starting with 100 different guesses across a wide range of values (x-axis) for each of the six parameters to be optimized - the internalization rate constant, fraction recycled, and endosomal exit rate constant of the ADC-Ag complex (top row); and the internalization rate constant, fraction recycled, and endosomal exit rate constant of the antigen (bottom row) - we obtained optimized values that for lowest-cost fits were not dependent on initial guess for some of the parameters but were for others. **B-C**, The simulation results (lines) for the lowest-cost optimization demonstrates good fit to the experimental data (dots).

A

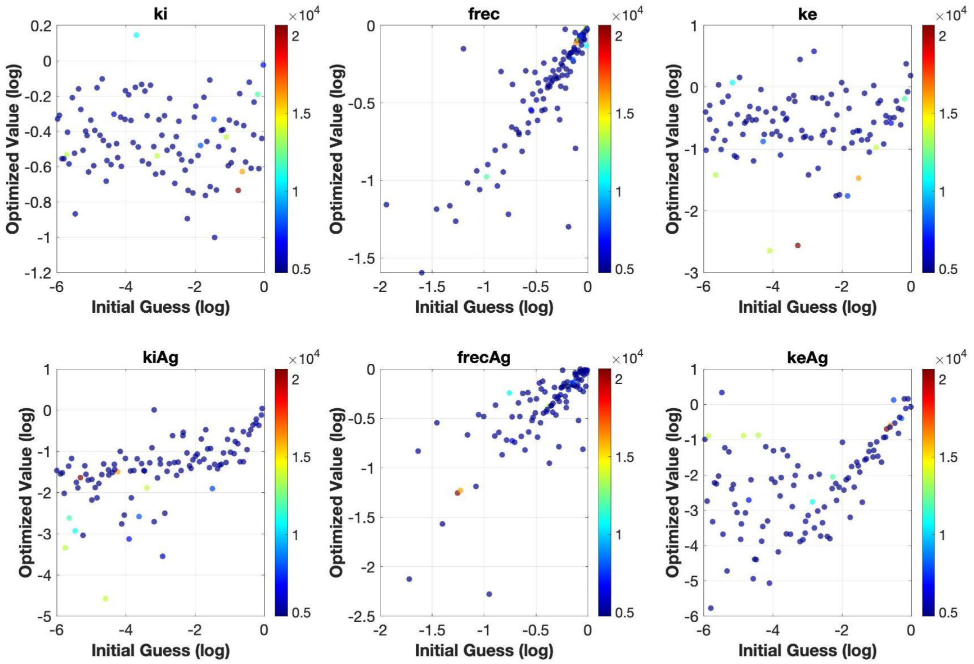

ki
frec
ke
kiAg
frecAg
keAg
Optimized Value (log)
Initial Guess (log)
×10^4

B

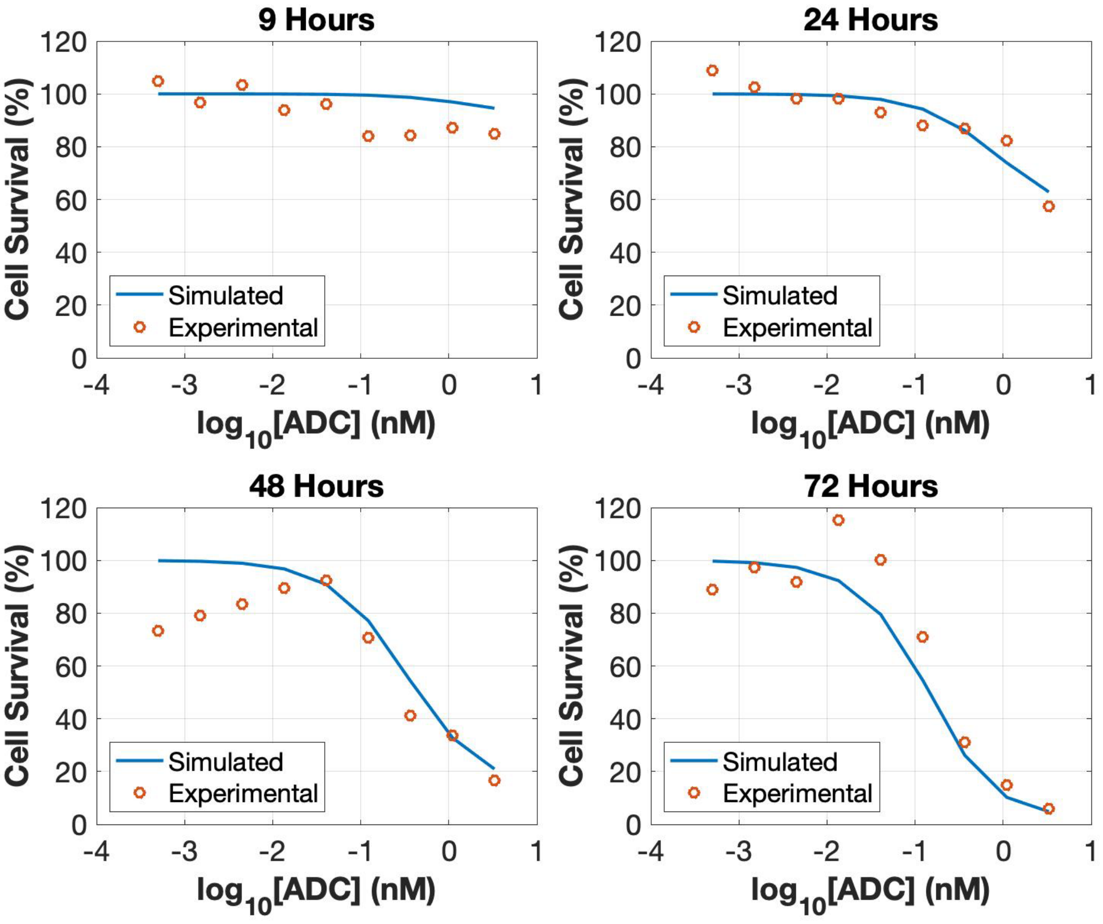
9 Hours
24 Hours
48 Hours
72 Hours
Cell Survival (%)
log10[ADC] (nM)
Simulated
Experimental
120
100
80
60
40
20
0
-4
-3
-2
-1
0
1

C

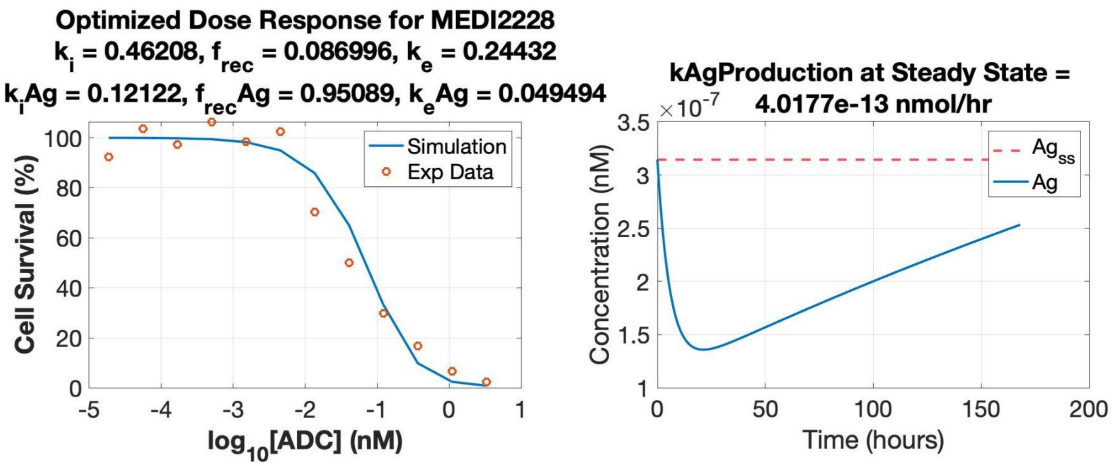
Optimized Dose Response for MEDI2228
ki = 0.46208, frec = 0.086996, ke = 0.24432
kiAg = 0.12122, frecAg = 0.95089, keAg = 0.049494
Simulation
Exp Data
Cell Survival (%)
log10[ADC] (nM)
kAgProduction at Steady State =
4.0177e-13 nmol/hr
×10-7
Agss
Ag
Concentration (nM)
Time (hours)

**Figure S6. Optimization of 3 linked receptor-mediated uptake parameters to ADC cytotoxicity data.** This includes nested optimization of antigen production rate to steady state levels in the absence of ADC. **A**, Starting with 100 different guesses across a wide range of values (x-axis) for each of the three parameters to be optimized - the internalization rate constant, fraction recycled, and endosomal exit rate constant for the ADC-Ag complex, with these parameters assumed to also apply to the antigen - we obtained optimized values that for lowest-cost fits were mostly not dependent on initial guess and demonstrated consistency/were well constrained. **B-C**, The simulation results (lines) for the lowest-cost optimization demonstrates good fit to the experimental data (dots).

**A**

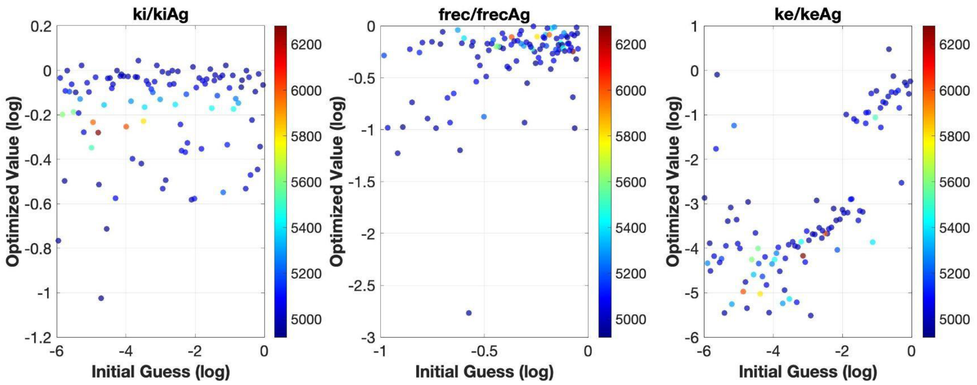
ki/kiAg
frec/frecAg
ke/keAg
Optimized Value (log)
Initial Guess (log)
0.2
0
-0.2
-0.4
-0.6
-0.8
-1
-1.2
-6
-4
-2
-0.5
-1.5
-2.5
-3
6200
6000
5800
5600
5400
5200
5000
1

**B**

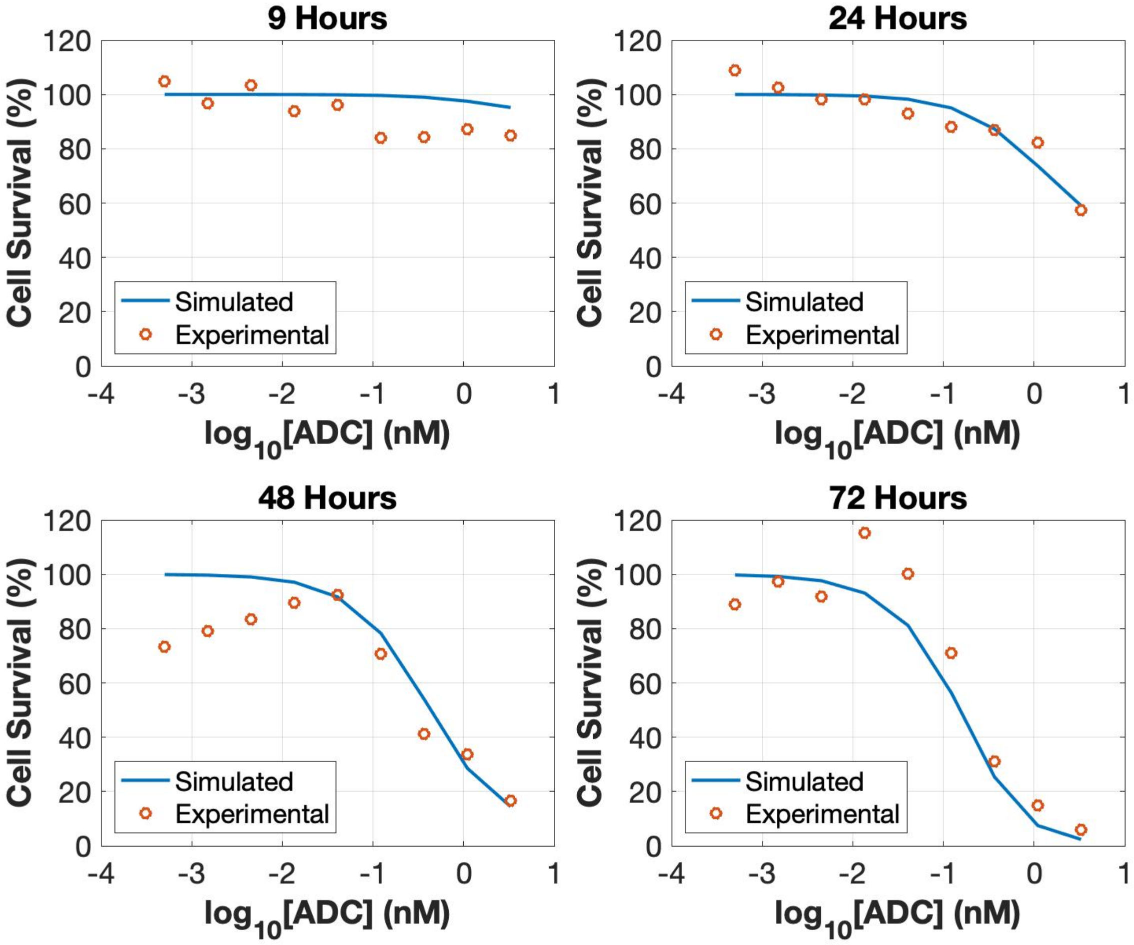
9 Hours
24 Hours
48 Hours
72 Hours
Cell Survival (%)
log10[ADC] (nM)
Simulated
Experimental
120
100
80
60
40
20
0
-4
-3
-2
-1
1

C

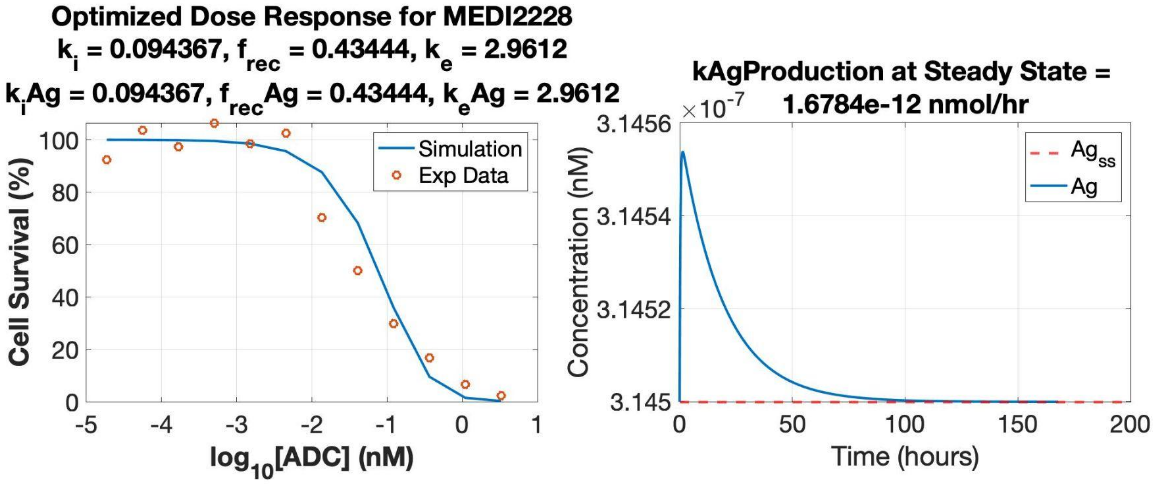
Optimized Dose Response for MEDI2228
k_i = 0.094367, f_rec = 0.43444, k_e = 2.9612
k_iAg = 0.094367, f_recAg = 0.43444, k_eAg = 2.9612
Simulation
Exp Data
Cell Survival (%)
log10[ADC] (nM)
kAgProduction at Steady State =
1.6784e-12 nmol/hr
×10^-7
Ag_ss
Ag
Concentration (nM)
Time (hours)

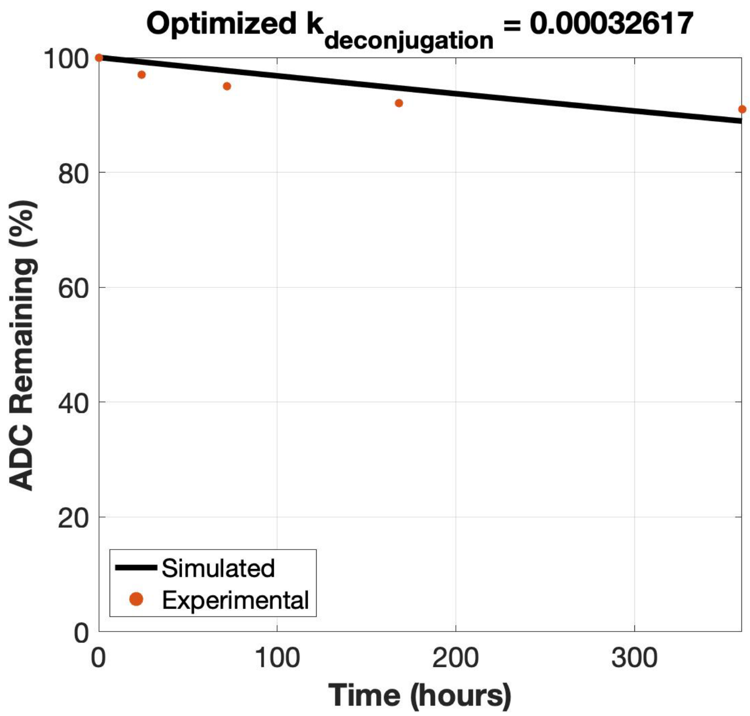


**Figure S7. Optimization of extracellular deconjugation rate constant to serum stability data.**
We optimized the rate constant to experimental data (dots) of percent ADC remaining over time, for ADC incubated at 200 ug/mL in mouse serum measured at 0, 1, 3, 7, and 15 days.

**Figure S8. Extracellular media volumes impact ADC efficacy and bystander effect.**
Area under the curve (AUC) of extracellular and intracellular warhead for ADCs with noncleavable linkers (**A**) and cleavable linkers (**B**). These are the AUC values for the full curves shown in Figure 8A and 8B respectively. Higher volumes (columns on the right) represent cell culture conditions, whereas lower volumes (columns on the left) more closely resemble physiologic conditions. As extracellular volumes decrease, the intracellular warhead source is more likely to come from influx than from higher volumes. At these lower volumes, a substantial amount of intracellular warhead comes from influx from other cells, suggesting that bystander effects may still play a significant role in ADCs with noncleavable linkers.

**A**

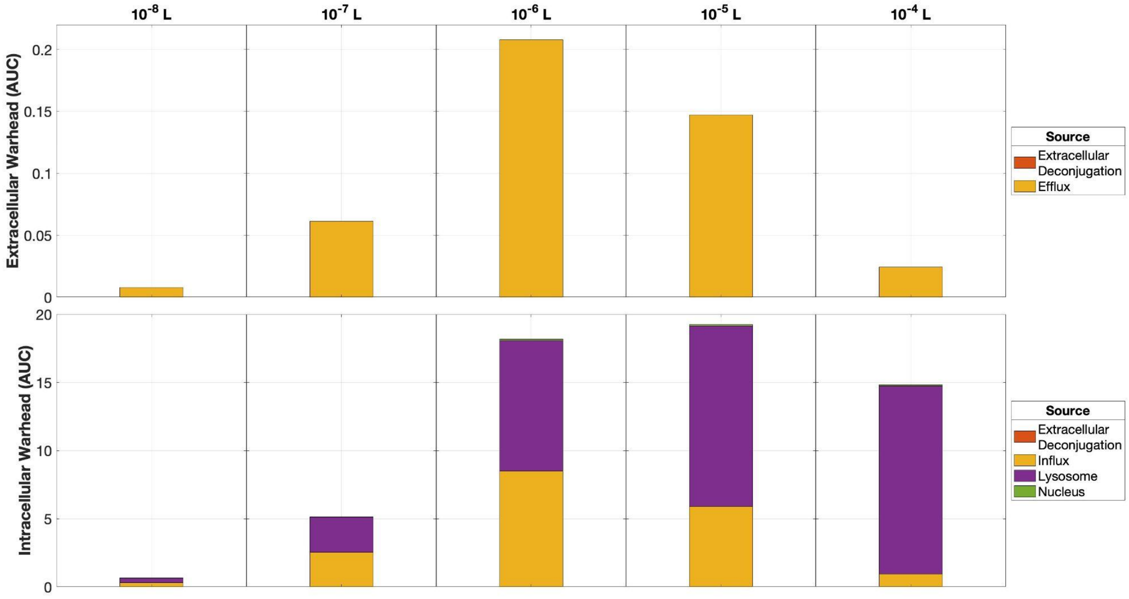
10^-8 L
10^-7 L
10^-6 L
10^-5 L
10^-4 L
Extracellular Warhead (AUC)
Intracellular Warhead (AUC)
0
0.05
0.1
0.15
0.2
0
5
10
15
20
Source
Extracellular Deconjugation
Efflux
Source
Extracellular Deconjugation
Influx
Lysosome
Nucleus

**B**

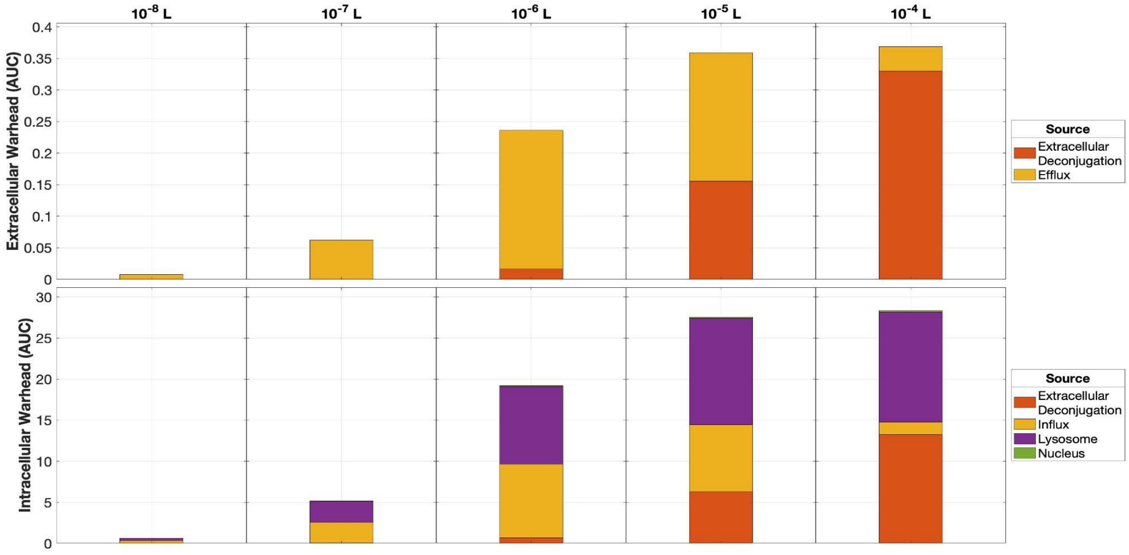
10^-8 L
10^-7 L
10^-6 L
10^-5 L
10^-4 L
Extracellular Warhead (AUC)
Intracellular Warhead (AUC)
0
0.05
0.1
0.15
0.2
0.25
0.3
0.35
0.4
0
5
10
15
20
25
30
Source
Extracellular Deconjugation
Efflux
Source
Extracellular Deconjugation
Influx
Lysosome
Nucleus

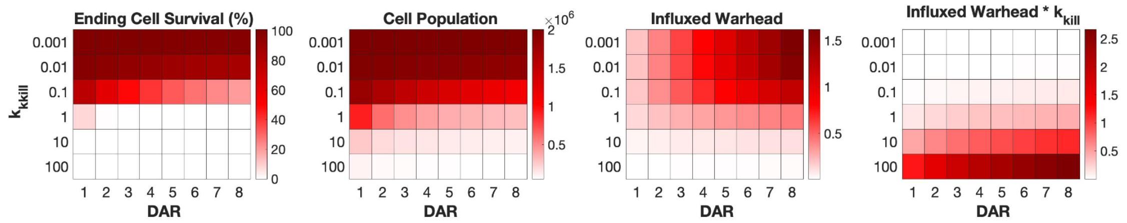


**Figure S9. Varying ADC design parameters (DAR and warhead potency, $k_{kill}$) to examine efficacy and toxicity outputs, including ending cell survival.**

Beyond the outputs of cell population, influxed warhead, and influxed warhead times warhead potency (from Figure S9), the ending cell survival can also be used as a metric for *in vitro* efficacy and shows a large decrease between $k_{kill}$ values of 0.1 and 1.

### Supplemental Methods

**Supplemental Methods S1. Computational Model Equations.** This is the system of coupled, nonlinear, ordinary differential equations we use to describe ADC, antibody, and warhead dynamics, interactions, and other processes in the media, on the cell surface, and in various locations in the cells.

**<u>Cell Surface / Extracellular Space</u>**

**[ADC] (nM)**: extracellular concentration of antibody-drug conjugate
Rate of change = infusion
+ (– ADC-Ag binding + unbinding – pinocytosis + FcRN recycling)*cell density
– deconjugation – ADC-solubleAg binding + unbinding

$$\frac{d[ADC]}{dt} = \frac{q}{V_{media}} + [Cells] * \begin{pmatrix} -k_{on,Ag} * [ADC] * [Ag] + k_{off,Ag} * [ADC \cdot Ag] \\ -\left(\frac{V_{endo}}{V_{media}}\right) * k_{i,ADC} * ([ADC] - [ADC_{endo}]) \\ +\left(\frac{V_{endo}}{V_{media}}\right) * k_{rec,ADC} * [ADC \cdot FcRn_{endo}] \end{pmatrix} - k_{deg} * [ADC] - k_{on,Ag_s} * [ADC] * [Ag_s] + k_{off,Ag_s} * [ADC \cdot Ag_s]$$

*Note: units of Vmedia = L; units of Vendo = L/cell; units of [Cells] = number (per well)*

**[Ag] (nM/cell)**: cell surface level of antigen
Rate of change = production – ADC-Ag binding + unbinding
– Ab-Ag binding + unbinding
– internalization + recycling

$$\frac{d[Ag]}{dt} = \frac{k_{AgProduction}}{V_{media}} - k_{on,Ag} * [ADC] * [Ag] + k_{off,Ag} * [ADC \cdot Ag] - k_{on,Ag} * [Ab] * [Ag] + k_{off,Ag} * [Ab \cdot Ag] - k_{i,Ag} * [Ag] + \left(\frac{V_{endo}}{V_{media}}\right) * f_{rec,Ag} * k_{e,Ag} * [Ag_{endo}]$$

**[ADC.Ag] (nM/cell)**: cell surface level of ADC-bound antigen
Rate of change = + ADC-Ag binding – unbinding
– internalization + recycling

$$\frac{d[ADC \cdot Ag]}{dt} = k_{on,Ag} * [ADC] * [Ag] - k_{off,Ag} * [ADC \cdot Ag] - k_i * [ADC \cdot Ag] + \left(\frac{V_{endo}}{V_{media}}\right) * f_{rec} * k_e * [ADC \cdot Ag_{endo}]$$

**[Ags] (nM)**: extracellular concentration of antigen
Rate of change = – ADC-Ags binding + unbinding
– Ab-Ags binding + unbinding

$$\frac{d[Ag_s]}{dt} = -k_{on,Ag_s} * [ADC] * [Ag_s] + k_{off,Ag_s} * [ADC \cdot Ag_s] - k_{on,Ag_s} * [Ab] * [Ag_s] + k_{off,Ag_s} * [Ab \cdot Ag_s]$$

**[ADC.Ags] (nM)**: extracellular concentration of ADC-bound antigen

Rate of change = + ADC-Ags binding – unbinding
– Ab-Ags binding + unbinding

$$\frac{d[ADC \cdot Ag_s]}{dt} = k_{on,Ag_s} * [ADC] * [Ag_s] - k_{off,Ag_s} * [ADC \cdot Ag_s]$$

**[Ab] (nM)**: extracellular concentration of antibody (without warhead)

Rate of change =
+ (– Ab-Ag binding + unbinding – pinocytosis + FcRN recycling)*cell density
+ generation from ADC deconjugation – Ab-solubleAg binding + unbinding

$$\frac{d[Ab]}{dt} = [Cells] * \begin{pmatrix} -k_{on,Ag} * [Ab] * [Ag] + k_{off,Ag} * [Ab \cdot Ag] \\ -\left(\frac{V_{endo}}{V_{media}}\right) * k_{i,ADC} * ([Ab] - [Ab_{endo}]) \\ +\left(\frac{V_{endo}}{V_{media}}\right) * k_{rec,ADC} * [Ab \cdot FcRn_{endo}] \end{pmatrix} + k_{deg} * [ADC] - k_{on,Ag_s} * [Ab] * [Ag_s] + k_{off,Ag_s} * [Ab \cdot Ag_s]$$

**[Ab.Ag] (nM/cell)**: cell surface level of Ab-bound antigen

Rate of change = + Ab-Ag binding – unbinding
– internalization + recycling

$$\frac{d[Ab \cdot Ag]}{dt} = k_{on,Ag} * [Ab] * [Ag] - k_{off,Ag} * [Ab \cdot Ag] - k_i * [Ab \cdot Ag] + \left(\frac{V_{endo}}{V_{media}}\right) * f_{rec} * k_e * [Ab \cdot Ag_{endo}]$$

**[Ab.Ags] (nM)**: extracellular concentration of Ab-bound soluble antigen

Rate of change = + Ab-Ags binding – unbinding

$$\frac{d[Ab \cdot Ag_s]}{dt} = k_{on,Ag_s} * [Ab] * [Ag_s] - k_{off,Ag_s} * [Ab \cdot Ag_s]$$

**Endosomal / Lysosomal Space**

**[ADCendo] (nM)**: endosomal concentration of ADC

Rate of change = + pinocytosis
– ADC-Ag binding + unbinding
– ADC-FcRN binding + unbinding
– exit rate (to lysosome)

$$\frac{d[ADC_{endo}]}{dt} = k_{i,ADC} * ([ADC] - [ADC_{endo}])$$
$$- k_{on,Ag} * [ADC_{endo}] * [Ag_{endo}] + k_{off,Ag} * [ADC \cdot Ag_{endo}]$$
$$- k_{on,FcRn} * [ADC_{endo}] * [FcRn_{endo}] + k_{off,FcRn} * [ADC \cdot FcRn_{endo}]$$
$$- k_{e,ADC} * [ADC_{endo}]$$

**[Agendo] (nM)**: endosomal concentration of antigen

Rate of change = + internalization
– ADC-Ag binding + unbinding
– Ab-Ag binding + unbinding
– recycling – degradation

$$\frac{d[Ag_{endo}]}{dt} = \left(\frac{V_{media}}{V_{endo}}\right) * k_{i,Ag} * [Ag]$$
$$- k_{on,Ag} * [ADC_{endo}] * [Ag_{endo}] + k_{off,Ag} * [ADC \cdot Ag_{endo}]$$
$$- k_{on,Ag} * [Ab_{endo}] * [Ag_{endo}] + k_{off,Ag} * [Ab \cdot Ag_{endo}]$$
$$- f_{rec,Ag} * k_{e,Ag} * [Ag_{endo}] - \left(1 - f_{rec,Ag}\right) * k_{e,Ag} * [Ag_{endo}]$$

**[ADC.Agendo] (nM)**: endosomal concentration of ADC-bound antigen

Rate of change = + internalization
+ ADC-Ag binding – unbinding
– recycling – degradation

$$\frac{d[ADC \cdot Ag_{endo}]}{dt} = \left(\frac{V_{media}}{V_{endo}}\right) * k_i * [ADC \cdot Ag]$$
$$+ k_{on,Ag} * [ADC_{endo}] * [Ag_{endo}] - k_{off,Ag} * [ADC \cdot Ag_{endo}]$$
$$- f_{rec} * k_e * [ADC \cdot Ag_{endo}] - (1 - f_{rec}) * k_e * [ADC \cdot Ag_{endo}]$$

**[Abendo] (nM)**: endosomal concentration of antibody (without warhead)

Rate of change = + pinocytosis
– Ab-Ag binding + unbinding
– Ab-FcRN binding + unbinding
– exit rate (to lysosome)

$$\frac{d[Ab_{endo}]}{dt} = k_{i,ADC} * ([Ab] - [Ab_{endo}])$$
$$- k_{on,Ag} * [Ab_{endo}] * [Ag_{endo}] + k_{off,Ag} * [Ab \cdot Ag_{endo}]$$
$$- k_{on,FcRn} * [Ab_{endo}] * [FcRn_{endo}] + k_{off,FcRn} * [Ab \cdot FcRn_{endo}]$$
$$- k_{e,ADC} * [Ab_{endo}]$$

**[Ab.Agendo] (nM)**: endosomal concentration of Ab-bound antigen

Rate of change = + internalization
+ Ab-Ag binding – unbinding
– recycling – degradation

$$\frac{d[Ab \cdot Ag_{endo}]}{dt} = \left(\frac{V_{media}}{V_{endo}}\right) * k_i * [Ab \cdot Ag]$$
$$+ k_{on,Ag} * [Ab_{endo}] * [Ag_{endo}] - k_{off,Ag} * [Ab \cdot Ag_{endo}]$$
$$- f_{rec} * k_e * [Ab \cdot Ag_{endo}] - (1 - f_{rec}) * k_e * [Ab \cdot Ag_{endo}]$$

**[FcRNendo] (nM)**: endosomal concentration of FcRN
Rate of change = – ADC-FcRN binding + unbinding
– Ab-FcRN binding + unbinding
+ recycling recovery

$$\frac{d[FcRn_{endo}]}{dt} = -k_{on,FcRn} * [ADC_{endo}] * [FcRn_{endo}] + k_{off,FcRn} * [ADC \cdot FcRn_{endo}]$$
$$- k_{on,FcRn} * [Ab_{endo}] * [FcRn_{endo}] + k_{off,FcRn} * [Ab \cdot FcRn_{endo}]$$
$$+ k_{rec,ADC} * [ADC \cdot FcRn_{endo}] + k_{rec,ADC} * [Ab \cdot FcRn_{endo}]$$

**[ADC.FcRNendo] (nM)**: endosomal concentration of ADC-bound FcRN
Rate of change = – ADC-FcRN binding + unbinding
– recycling

$$\frac{d[ADC \cdot FcRn_{endo}]}{dt} = k_{on,FcRn} * [ADC_{endo}] * [FcRn_{endo}] - k_{off,FcRn} * [ADC \cdot FcRn_{endo}]$$
$$- k_{rec,ADC} * [ADC \cdot FcRn_{endo}]$$

**[Ab.FcRNendo] (nM)**: endosomal concentration of Ab-bound FcRN
Rate of change = – Ab-FcRN binding + unbinding
– recycling

$$\frac{d[Ab \cdot FcRn_{endo}]}{dt} = k_{on,FcRn} * [Ab_{endo}] * [FcRn_{endo}] - k_{off,FcRn} * [Ab \cdot FcRn_{endo}]$$
$$- k_{rec,ADC} * [Ab \cdot FcRn_{endo}]$$

**[Xlys] (nM)**: lysosomal concentration of X (X = ADC, Ag, and complexes)
Rate of change = + degradation – loss

$$\frac{d[ADC_{lys}]}{dt} = k_{e,ADC} * [ADC_{endo}] - [ADC_{lys}]$$
$$\frac{d[Ag_{lys}]}{dt} = (1 - f_{rec,Ag}) * k_{e,Ag} * [Ag_{endo}] - [Ag_{lys}]$$
$$\frac{d[ADC \cdot Ag_{lys}]}{dt} = (1 - f_{rec}) * k_e * [ADC \cdot Ag_{endo}] - [ADC \cdot Ag_{lys}]$$
$$\frac{d[Ab_{lys}]}{dt} = k_{e,ADC} * [Ab_{endo}] - [Ab_{lys}]$$
$$\frac{d[Ab \cdot Ag_{lys}]}{dt} = (1 - f_{rec}) * k_e * [Ab \cdot Ag_{endo}] - [Ab \cdot Ag_{lys}]$$

**<u>Clearance</u>**

**[ADCcl] (nM)**: total degraded antibody (accumulates over time)

Rate of change = degradation of ADC-containing components

$$\frac{d[ADC_{cl}]}{dt} = \left[ADC_{lys}\right] + \left[ADC \cdot Ag_{lys}\right]$$

**[Agcl] (nM)**: total degraded antigen (accumulates over time)
Rate of change = degradation of Ag-containing components

$$\frac{d[Ag_{cl}]}{dt} = \left[Ag_{lys}\right] + \left[ADC \cdot Ag_{lys}\right] + \left[Ab \cdot Ag_{lys}\right]$$

**[Abcl] (nM)**: total degraded antibody (accumulates over time)
Rate of change = degradation of Ab-containing components

$$\frac{d[Ab_{cl}]}{dt} = \left[Ab_{lys}\right] + \left[Ab \cdot Ag_{lys}\right]$$

**<u>Cytoplasm</u>**

**[Win] (nM)**: intracellular warhead
Rate of change = release by degraded lysosomal ADC
– efflux of warhead + influx of warhead
– nuclear entry + return from nucleus

$$\frac{d[W_{in}]}{dt} = \left(\frac{V_{endo}}{V_{cytoplasm}}\right) * DAR * \left(\left[ADC_{lys}\right] + \left[ADC \cdot Ag_{lys}\right]\right)$$
$$- k_{eff} * [W_{in}] + \left(\frac{V_{media}}{V_{cytoplasm}}\right) * \left(\frac{V_{cytoplasm}}{V_{media}}\right) * k_{inf} * [W_{ex}]$$
$$- k_{in,nuc} * [W_{in}] + \left(\frac{V_{nuc}}{V_{cytoplasm}}\right) * k_{out,nuc} * [W_n]$$

**<u>Intracellular Warhead Tracking</u>**

**[Win,lys] (nM)**: intracellular warhead most recently released by lysosomal degradation
Rate of change = release by degraded lysosomal ADC
– efflux of warhead – nuclear entry

$$\frac{d[W_{in,lys}]}{dt} = \left(\frac{V_{endo}}{V_{cytoplasm}}\right) * DAR * \left(\left[ADC_{lys}\right] + \left[ADC \cdot Ag_{lys}\right]\right) - \left(k_{eff} + k_{in,nuc}\right) * \left[W_{in,lys}\right]$$

**[Win,deg] (nM)**: intracellular warhead most recently derived by influx of extracellular deconjugated warhead
Rate of change = influx by degraded lysosomal ADC
– efflux of warhead – nuclear entry

$$\frac{d[W_{in,deg}]}{dt} = \left(\frac{V_{media}}{V_{cytoplasm}}\right) * \left(\frac{V_{cytoplasm}}{V_{media}}\right) * k_{inf} * [W_{ex,deg}] - (k_{eff} + k_{in,nuc}) * [W_{in,deg}]$$

*Note: typically movement between two compartments can be modeled using a dilution effect; however, the size of the extracellular media is so much higher than the cytoplasm that this would quickly result in an overwhelming spike in intracellular concentration. Therefore, we include a reciprocal correction to reflect that the influx is typically sampling from media volume near the cells, and the influx rate constant value will take care of the rest.*

**[Win,influx] (nM)**: intracellular warhead most recently derived by influx of extracellular deconjugated warhead

Rate of change = influx of previously effluxed warhead
– efflux of warhead – nuclear entry

$$\frac{d[W_{in,influx}]}{dt} = \left(\frac{V_{media}}{V_{cytoplasm}}\right) * \left(\frac{V_{cytoplasm}}{V_{media}}\right) * k_{inf} * [W_{ex,efflux}] - (k_{eff} + k_{in,nuc}) * [W_{in,influx}]$$

**[Win,nuc] (nM)**: intracellular warhead most recently derived by export from the nucleus

Rate of change = export of warhead from the nucleus
– efflux of warhead – nuclear entry

$$\frac{d[W_{in,nuc}]}{dt} = \left(\frac{V_{nuc}}{V_{cytoplasm}}\right) * k_{out,nuc} * [W_n] - (k_{eff} + k_{in,nuc}) * [W_{in,nuc}]$$

### Extracellular Space

**[Wex] (nM)**: extracellular warhead

Rate of change = influx of previously effluxed warhead
+ (efflux of warhead from cells – influx into cells)*cell density

$$\frac{d[W_{ex}]}{dt} = DAR * k_{deg} * [ADC] + [Cells] * \left(\frac{V_{cytoplasm}}{V_{media}}\right) * (k_{eff} * [W_{in}] - k_{inf} * [W_{ex}])$$

*Note: units of Vmedia = L; units of Vcytoplasm = L/cell; units of [Cells] = number (per well)*

### Extracellular Warhead Tracking

**[Wex,deg] (nM)**: extracellular warhead most recently derived by deconjugated warhead

Rate of change = deconjugation from ADC
– influx into cells

$$\frac{d[W_{ex,deg}]}{dt} = DAR * k_{deg} * [ADC] - [Cells] * \left(\frac{V_{cytoplasm}}{V_{media}}\right) * k_{inf} * [W_{ex,deg}]$$

**[Wex,efflux] (nM)**: extracellular warhead most recently derived by efflux from cells

Rate of change = efflux from cells – influx into cells

$$\frac{d[W_{ex,efflux}]}{dt} = [Cells] * \left(\frac{V_{cytoplasm}}{V_{media}}\right) * \left(k_{eff} * [W_{in}] - k_{inf} * [W_{ex,efflux}]\right)$$

**<u>Nucleus</u>**

**[Wn] (nM)**: nuclear warhead

Rate of change = – warhead-DNA binding + unbinding
+ nuclear entry – export from nucleus

$$\frac{d[W_n]}{dt} = -k_{on,DNA} * [W_n] * [DNA] + k_{off,DNA} * [W_n \cdot DNA] + \left(\frac{V_{cytoplasm}}{V_{nuc}}\right) * k_{in,nuc} * [W_{in}] - k_{out,nuc} * [W_n]$$

**[DNA] (nM)**: density of warhead-binding DNA sites

Rate of change = – warhead-DNA binding + unbinding

$$\frac{d[DNA]}{dt} = -k_{on,DNA} * [W_n] * [DNA] + k_{off,DNA} * [W_n \cdot DNA]$$

**[Wn.DNA] (nM)**: warhead-bound DNA sites

Rate of change = + warhead-DNA binding – unbinding

$$\frac{d[W_n \cdot DNA]}{dt} = k_{on,DNA} * [W_n] * [DNA] - k_{off,DNA} * [W_n \cdot DNA]$$

**<u>Crosslink Formation</u>**

$$f_{crosslinked} = \frac{1}{1 + \left(\frac{K_A}{[W_n \cdot DNA]}\right)^n}$$

**<u>Cell Growth and Cell Killing</u>**

Rate of change = cell growth – crosslink-induced cell death

$$\frac{d[Cells]}{dt} = k_g * [Cells] - f_{crosslinked} * k_{kill} * [Cells]$$
